# Host and microbial factors influence bacterial colonization of the honey bee gut

**DOI:** 10.64898/2025.12.04.691542

**Authors:** Silvia Moriano-Gutierrez, Aurélien Pirat, Audam Chhun, Aiswarya Prasad, Alexandra Szigeti, Méline Garcia, Lucie Kesner, Florent Mazel, Philipp Engel

## Abstract

The guts of many animals are colonized by host-specific microbes, yet the extent to which host filtering (host-derived constraints) shapes microbial colonization and host specificity remains poorly understood. Here, we used gnotobiotic honey bees (*Apis mellifera*) as a model system to systematically assess the colonization potential of a phylogenetically and ecologically diverse panel of 56 bacterial strains, spanning native symbionts, opportunistic bee-associated taxa, gut microbes from other bee species, and non-bee environmental isolates. Bacterial load and colonization frequency were quantified by strain-specific qPCR seven days post-inoculation, in monocolonization and in the presence of a synthetic community composed of native honeybee core bacteria. Bacterial load was highest for native strains and declined with increasing phylogenetic distance from native symbionts. Co-colonization with the synthetic community reduces load across all groups, but native strains were least affected. Across strains, completeness of KEGG metabolic pathways correlated with bacterial load in some ecological groups; however, metabolic capacity alone did not fully explain colonization patterns, either in monocolonization or under competitive conditions. A key finding was that *in vitro* sensitivity to antimicrobial peptide (AMPs; apidaecin, abaecin, defensins, hymenoptaecin) varied widely among strains and was highest in closely related bee-associated bacteria. Notably, even highly successful colonizers such as *Gilliamella* and *Snodgrassella* were AMP-sensitive. AMP sensitivity showed a negative correlation with bacterial load, but not with the frequency of host colonization. These findings suggest that AMPs modulate symbiont abundance rather than acting as strict barriers to colonization. Overall, our results reveal that host filtering in the bee gut is multifaceted, integrating immune-mediated barriers, microbial traits, and competitive interactions.

## INTRODUCTION

The animal gut microbiome fulfills key functions for the host in nutrition, development, defense and even behaviour^1–7^. While gut symbionts are horizontally transmitted, they usually exhibit a high degree of host specificity^8–11^. Understanding the mechanisms that underlie microbiota assembly and symbiont specificity remains a central question in microbiome research. Multiple factors, including host filtering (host-derived constraints on microbial establishment and/or abundance), microbe-microbe interactions (competition/facilitation), and dispersal limitation, are known to contribute^8,9,12,13^, but their relative influence, particularly across microbial strains with different evolutionary histories, is still not fully understood. Evidence from other host systems shows that colonization potential is not determined by isolation source alone: microbes from diverse environments can establish in the gut when they possess traits suited to that habitat^14^. This raises the question of whether similar trait-based mechanisms explain colonization by non-native bacteria, and how they interact with host-specific constraints and resident microbiota. The assembly of animal gut microbiota is governed by a complex interplay of host and microbial factors that collectively shape which microbes establish and persist. From the host side, immune responses, gut physiology, and nutrient availability act as selective filters that can promote native symbionts while excluding non-native taxa^8^. However, it remains unclear whether native and non-native taxa differ enough for the honey bee’s innate inmmune system to differentially detect and constrain them, including those from closely related hosts. From the microbial perspective, colonization requires overcoming several ecological and physiological challenges: entering the host, reaching and replicating in specific gut niches, resisting immune defenses, and interacting with competing microbes^15^. Traits such as antimicrobial peptide (AMP) resistance, metabolic compatibility and immune evasion all contribute to whether a strain can establish in the gut^16–22^. Additionally, ecological interactions between microbes, including competition (e.g., nutrient exclusion, niche preemption) and facilitation (e.g., cross-feeding, immune modulation), can shape community outcomes^19,23–26^. These interactions are further constrained by behavioral and ecological barriers of hosts, such as dietary habits or restricted transmission routes, which limit dispersal and reinforce host-specific assemblages^10,27,28^.

Honey bees (*Apis mellifera*) serve as a powerful model organism for investigating factors underlying host-microbe specificity. As social corbiculate bees, they harbor a simple yet highly specialized gut microbiota that can be readily studied under controlled laboratory conditions^29^. This community comprises approximately 15-20 bacterial species across a few dominant genera^30–32^. Adult worker guts are consistently dominated by five “core” genera: *Gilliamella*, *Snodgrassella*, *Lactobacillus, Bombilactobacillus* and *Bifidobacterium*. Additional bee-associated genera, such as *Frischella*, *Bartonella*, and *Commensalibacter,* are frequently detected but are generally considered non-core, with prevalence varying among individuals and colonies^33^. Opportunistic colonizers, such as environmentally acquired commensals or pathogens (e.g. *Serratia* or *Hafnia*) are also occasionally detected but occur relatively rarely^34^. This characteristic gut microbiota is primarily acquired through social transmission shortly after adult emergence, via contact with nest-mates and hive material^35,36^. Although the gut microbiota of other social bees, such as other *Apis* species, bumble bees, and stingless bees, often include members of the same genera, these bacteria are typically represented by host-specific species or strains. Moreover, these symbionts are rarely found outside of the gut of bees, suggesting a high degree of host specificity^33^. However, the ecological processes and molecular mechanisms underlying the observed specificity are not fully understood. Experimental studies with gnotobiotic bees have demonstrated that bacteria of *Snodgrassella, Gilliamella,* and *Lactobacillus* generally exhibit greater colonization success in their native bee host than in other bee species^30,32^. Bacterial colonization of the bee gut activates host immune responses, including AMP production and modulation of key signaling pathways such as the Toll and Imd pathways^37–39^. In one specific case, it was shown that a non-native, but not a native strain, of the genus *Gilliamella* triggers the production of reactive oxygen species thereby partially hindering gut colonization^39^. To date, however, most studies have examined only a small number of strains, sometimes belonging to different species and often limited to a single “native vs. non-native” comparison. As a result, it remains unclear whether gut colonization is truly restricted to the recurrent core genera of social bees, or whether additional, underexplored lineages can also establish under certain conditions. Moreover, substantial variation in colonization efficiency, metabolic capacity, and immunogenicity is known even among closely related strains within core taxa^40,41^. Such intra- and interspecific diversity highlights the need for phylogenetically broad, strain-resolved experimental surveys that move beyond a few model isolates to broaden inference about the true breadth of colonization potential across ecologically defined bacterial groups.

In this study, we investigated the factors influencing bacterial colonization success in the honey bee gut using gnotobiotic bees. We assembled a diverse panel of 56 phylogenetically and ecologically distinct bacterial strains, grouped into four categories: (i) native symbionts belonging to the typical genera found in the honeybee gut microbiota, (ii) opportunistic colonizers of the honey bee gut, (iii) gut symbionts from other bee species, and (iv) non-bee environmental isolates. By evaluating colonization performance in both mono-association and community settings, and by characterizing AMP sensitivity, metabolic profiles, and interbacterial interactions, we identify bacterial traits and host factors that shape colonization outcomes. This integrated approach provides new insights into the complex and multifactorial nature of microbiota assembly and host specificity in animal symbioses.

## MATERIALS AND METHODS

### Generation of gnotobiotic bees

Late-stage pupae with pigmented eyes and light grey cuticle were removed from brood frames and incubated for 3 days at 35 °C and 75% humidity in sterile boxes, receiving sterile 1:1 sucrose solution. Upon emergence, bees were randomly assigned to one of four groups: microbiota-depleted (MD), colonized with a synthetic (or defined) community (SC) of native symbionts, monocolonized with individual strains (S), colonized with SC plus an additional strain (SC + S). Strains, detailed in Supplementary File 1, were grown, washed, standardized to OD₆₀₀ = 1.0, and stored in glycerol at −80 °C. The 14-member SC was assembled to represent the dominant worker gut symbiont genera of *A. mellifera* and was selected based on (i) assignment to the most prevalent genera present in the gut of *A. mellifera*, and (ii) reproducible colonization in gnotobiotic bees. The 14 strains comprising the SC are listed in Supplementary File 1 under category *Native*. For colonization, bees were fed 5 μL of bacterial suspension, of the glycerol stocks stored at −80 °C, in PBS:sucrose. S treatments received a single strain (OD₆₀₀ = 0.1); SC included 14 native strains mixed at an OD₆₀₀ = 0.1; SC + S combined SC with one added strain. MD bees received mock inoculum (1:1 PBS-sucrose solution). Guts were dissected seven days post-inoculation and stored at −80 °C. Seven days post-inoculation is commonly used in gnotobiotic honey bee assays because gut colonization and community structure stabilize within the first week of adulthood^42^. The experiment included three biological replicates using pupae from three different hives to account for across-colony variation.

### DNA extraction from honey bee gut tissue

DNA was extracted from honey bee gut samples using bead-beating followed by magnetic bead-based purification. Samples were kept on ice and processed alongside the negative controls. Each sample was lysed with ATL buffer containing Proteinase K (2 mg/mL) and mechanically disrupted using a bead-beater (6 m/s, 45 s). After incubation at 56 °C for 1 h with agitation, lysates were cleared by centrifugation, and DNA was purified using CleanNGS magnetic beads. DNA binding, ethanol washes, and elution were performed in a 96-well plate format. Final DNA eluates (40 μL) were stored at −20 °C for downstream analysis.

### RNA extraction from honey bee gut tissue

Total RNA was extracted from whole gut samples using the RNeasy PowerMicrobiome Kit (Qiagen), following the manufacturer’s instructions. Immediately after extraction, samples were treated with TURBO DNase (Thermo Fisher Scientific) to eliminate residual genomic DNA. RNA concentration for each sample was quantified using the Qubit RNA Broad Range (BR) Assay Kit (Invitrogen), according to the manufacturer’s protocol.

### Bacterial quantification by quantitative PCR (qPCR)

qPCR was performed on a QuantStudio™ 5 system (Applied Biosystems) in 96- or 364-well plates, using 10 μL reactions with 0.2 μM primers, 1× SYBR® Select Master Mix, and 1 μL DNA template. Thermal cycling included 50 °C for 2 min, 95 °C for 2 min, followed by 40 cycles of 95 °C for 15 s and 60 °C for 1 min. Each sample was run in triplicate. Total bacterial load, host gene expression, and strain-specific abundance were assessed using universal, species-specific, and host reference primers (see Supplementary File 2). Universal 16S rRNA primers were used only for MD and SC controls to assess background signal and total bacterial load, whereas strain-specific primers were used in inoculated treatments to quantify inoculated strain abundance and prevalence. Primer design followed Primer3plus; quantification methods are described in Kešnerová et al^43^.

### Host immune expression by qRT-PCR

For host immune gene expression, cDNA was synthesized from gut RNA using SMART MMLV reverse transcriptase with both Oligo(dT) and random hexamers. qRT-PCR followed MIQE guidelines, including no-RT and no-template controls. Standard curves ensured amplification efficiency, and reactions used 12.5 ng cDNA in triplicate. Melting curve analysis confirmed specificity. Gene expression was normalized to the *Apis mellifera* RPS18 reference gene. Details on all primer sequences and target genes are provided in Supplementary File 2.

### DNA isolation for genome sequencing

Genomic DNA for Illumina sequencing was extracted using the FastPure Bacterial DNA Isolation Mini Kit (Vazyme), following the manufacturer’s protocol for Gram-positive bacteria. Illumina sequencing libraries were prepared using the Nextera DNA Flex Library Prep Kit (Illumina), following the guidelines in the Illumina Reference Guide. Library indexing, dilution, and denaturation were performed according to the Illumina *Index Adapters Pooling Guide* and the *MiniSeq System Denature and Dilute Libraries Guide*. Final libraries were quantified using a double-stranded DNA fluorescent dye assay and sequenced on a MiniSeq high-output flow cell (150 bp paired-end reads).

### Genome assembly and functional annotation

Raw Illumina reads were quality-checked using FastQC v0.11.9^44^ and trimmed with Trimmomatic v0.39^45^ using the following parameters: PE-phred33 AllIllumina-PEadapters.fa:3:25:7 LEADING:9 TRAILING:9 SLIDINGWINDOW:4:15 MINLEN:60. Trimmed reads were assembled *de novo* using SPAdes v3.15.2 with the --careful option^46^. Genome completeness and contamination were evaluated using CheckM v1.0.13^47^(Supplementary File S7). Genomes were annotated with Prokka v1.13^48^. Open reading frames (ORFs) were identified on all assembled scaffolds of each sample using Prodigal^49^. ORFs marked partial or shorter than 300 bp were removed. The remaining ORFs were then annotated using DRAM^50^ to obtain KEGG ortholog (KO) identities.

### Phylogenetic analysis

A phylogenetic tree of the 56 bacterial species and strains used in colonization experiments was constructed using OrthoFinder v2.5.4. Orthologous gene groups were identified across all genomes, and species tree inference was performed using the STAG (Species Tree from All Genes) algorithm^51^, as implemented in OrthoFinder^52^. This approach integrates gene tree topologies to produce a robust consensus species tree.

### KEGG module annotation and metabolic profiling

Genomes were annotated and KEGG module completeness quantified per strain, generating a strain × module matrix (Supplementary File 7). Modules with ≥80% completeness were considered complete. For each strain, we calculated module richness and module-level completeness values for association testing. PCA was performed on the centered and scaled matrix. Module–abundance associations were tested with per-module mixed-effects models (log_median ∼ module + Category + Colonization_mode + (1 | genus)) and FDR correction. Module richness was tested against abundance with lme4::lmer(log_median_rank ∼ Number.of.modules + Category + (1 | genus)); when genus-level variance was ∼0, the corresponding fixed-effects model is also reported. Differences in module richness among categories were tested with number of modules ∼ Category + (1 | genus).

### Antimicrobial peptide resistance assays

To evaluate bacterial sensitivity to AMPs, resistance assays were conducted using a 12-point, two-fold serial dilution series of each AMP, ranging from 100 μg/mL to 0.0487 μg/mL. AMPs were synthesized by Shanghai Royobiotech Co., Ltd., with peptide sequences detailed in Supplementary File 6. A subset of strains was selected for AMP screening to represent each ecological category and to span the range of *in vivo* colonization outcomes; strains were additionally prioritized based on their ability to grow reproducibly and reach sufficient biomass under the in vitro assay conditions. Each bacterial strain was tested in triplicate across all peptide concentrations. Bacterial cultures were inoculated into 96-well microtiter plates containing AMP dilutions in 50% growth medium and incubated under standard conditions (Supplementary File 1). Bacterial growth was monitored by measuring optical density at 600 nm (OD₆₀₀) over time to assess biomass accumulation and peptide-induced growth inhibition. Growth inhibition was evaluated relative to untreated controls. Normalized area under the curve values were compared to a control mean of 1 using one-sample *t*-tests at each concentration and condition. *P*-values were adjusted for multiple testing using the FDR method. Growth was considered significantly inhibited at a given concentration if *q* < 0.05. A sensitivity score from 0 to 12 was assigned to each strain-peptide combination, representing the number of concentrations at which significant growth inhibition was detected. A score of 12 indicated growth inhibition even at the lowest concentration tested, while a score of 0 indicated no significant inhibition at any concentration.

### Statistical analyses

All statistical analyses were performed in R (version 4.2.2). Data are presented as mean ± standard deviation for approximately normally distributed data and as median with 95% confidence intervals otherwise. For parametric models, assumptions were checked on the model residuals. Normality of residuals was assessed using the Shapiro-Wilk test and visual inspection of Q-Q plots, and homoscedasticity was evaluated from residual-versus-fitted plots. Depending on the data distribution, statistical comparisons were performed using one-way ANOVA or Kruskal-Wallis tests. Proportion data were analyzed using chi-square tests for equality of proportions, with FDR correction for multiple comparisons where applicable. A p-value < 0.05 was considered statistically significant. Where applicable, p-values were adjusted for multiple comparisons. Sample size (n) refers to the number of independent biological replicates. Independent experimental replicates are indicated when applicable. Detailed information on statistical tests used is provided in the corresponding figure legends.

## RESULTS

### Bacterial strains vary in their ability to colonize the honey bee gut regardless of their isolation source

Using gnotobiotic bees, we screened 56 taxonomically and ecologically diverse bacterial strains for their ability to colonize the midgut and hindgut of the Western honey bee (Apis mellifera), either alone or in the presence of a synthetic community representing eight predominant gut genera. The strains belonged to 25 genera spanning Gamma- and Betaproteobacteria, Actinobacteria, and Firmicutes (Figure 1A) and fell into four ecological categories (Figure 1B, Supplementary File 1): (i) Native, isolated from A. mellifera and representing predominant gut genera; (ii) Opportunistic colonizers, isolated from A. mellifera but not belonging to predominant genera and typically detected at low abundance or prevalence; (iii) Non-native bee-associated, belonging to the same prevalent genera as native strains but isolated from other bee species and representing distinct bacterial species; and (iv) Other microbiomes, originating from non-bee environments such as soil, fish, and human or mouse guts. Seven days after inoculation, loads of all 56 strains were quantified by strain-specific qPCR (Figure 1A-C, Figure S1). Universal 16S rRNA qPCR of MD and SC controls (Figure S1) showed low background signal in MD bees and significantly higher bacterial loads in SC bees. Both the bacterial load per gut and the proportion of bees with levels exceeding the initial inoculum varied substantially across strains, ranging from 10^3^ to 10^10^ cells per bee (Figure 1C, Supplementary File 2) and from 0% to 100%, respectively (Figure S2).

**Figure 1.**
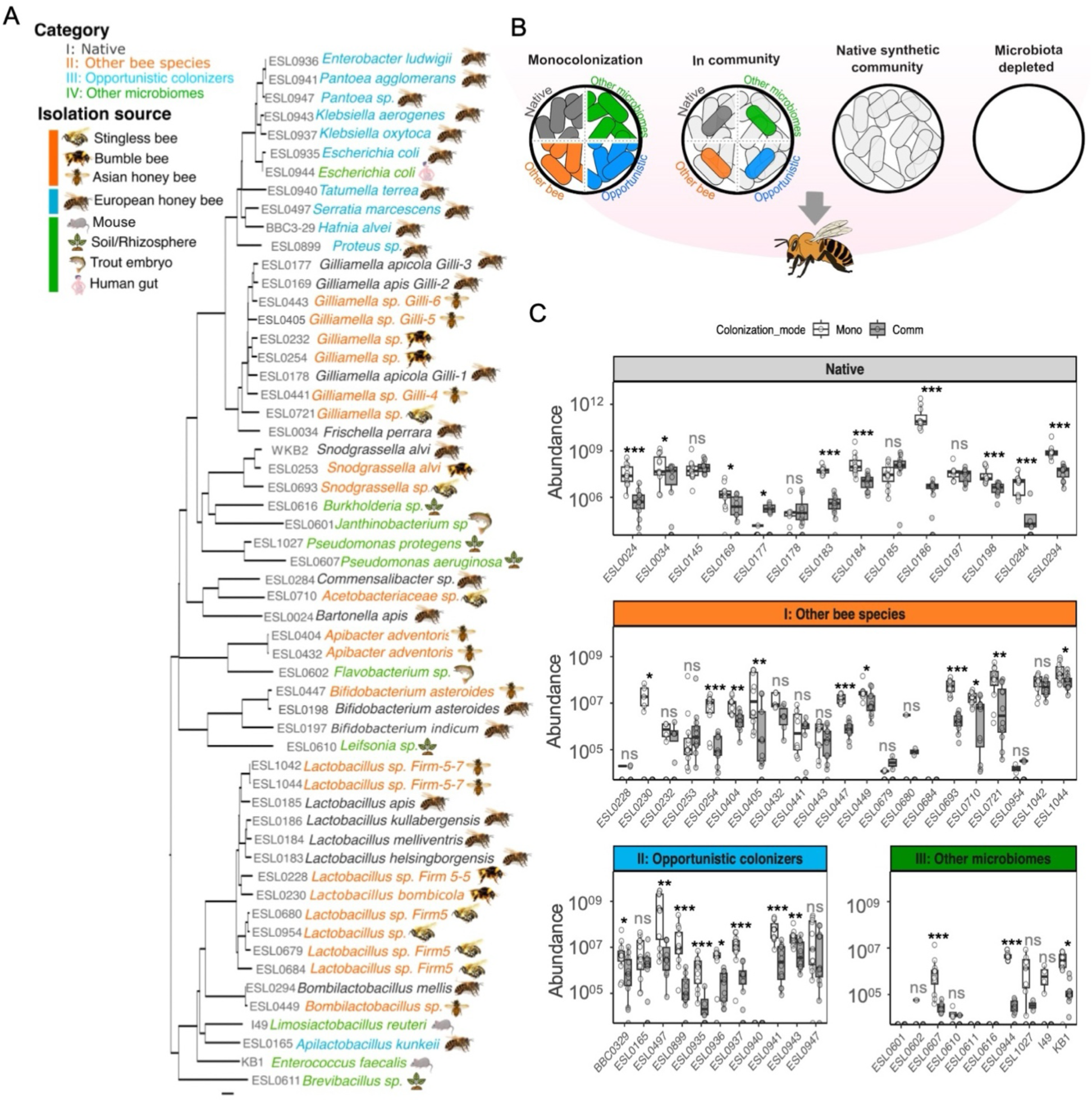
Bacterial colonization efficiency of the honey bee gut in monocolonization or in the presence of a native community. **A.** Phylogenetic tree of the 56 bacterial species and strains tested for colonization of the honey bee gut. The tree was inferred with OrthoFinder using the STAG algorithm^51^. Scale bar = 0.09 substitutions per site. **B.** Schematic outline of the *in vivo* experiment to estimate colonization levels of strains categorized as *Native* (n=14), *Other bee species* (n=21), *Opportunistic colonizers* (n=11) and *Other microbiomes* (n=10). For every colonization experiment a microbiota-depleted (MD) and a native synthetic community (SC) control was included (Figure S1). **C.** Absolute bacterial loads in the gut for all strains in monocolonization (Mono) or community (Comm), measured seven days post-inoculation. Group differences in population size were evaluated with a Kruskal-Wallis test (multiple groups), followed by pairwise Wilcoxon rank-sum tests with Benjamini-Hochberg FDR correction. The p-values were adjusted using FDR method. Significance labels shown as p<=0.05:*, p=<=0.01:**, p<=0.001:***, ns: not significant. Per strain 12 bees from three independent experimental replicates conducted with bees from different colonies were analyzed.

On average, the highest colonization loads were observed for strains derived from the native gut microbiota of *Apis mellifera*, followed by non-native bee-associated strains and opportunistic colonizers. In contrast, isolates originating from non-bee environments exhibited the lowest colonization levels (Figure 2A). Similarly, the proportion of colonization was significantly higher for native strains compared to those derived from other bee species (≈21% lower relative to native) or from other microbiomes (≈67% lower relative to native) (Figure 2B). Opportunistic colonizers of *Apis mellifera* reached a similar proportion of colonization success as native strains, though with a lower bacterial load. When introduced alongside a synthetic community representing the native microbiota, the differences in colonization patterns across the four ecological groups were comparable to those observed in the monocolonizations (Figure S2); however, both bacterial load and proportion of colonization were generally reduced, likely due to competitive interactions with the native community (Figure 1C, Figure 2C). Only one native strain, *Gilliamella* ESL0177, exhibited enhanced growth in the presence of the synthetic community compared to monocolonization. While this strain colonized poorly when introduced alone (Figure 1C, Figure S2), its bacterial load increased substantially in a community context, suggesting positive interactions with other native microbiota members facilitating its colonization

**Figure 2.**
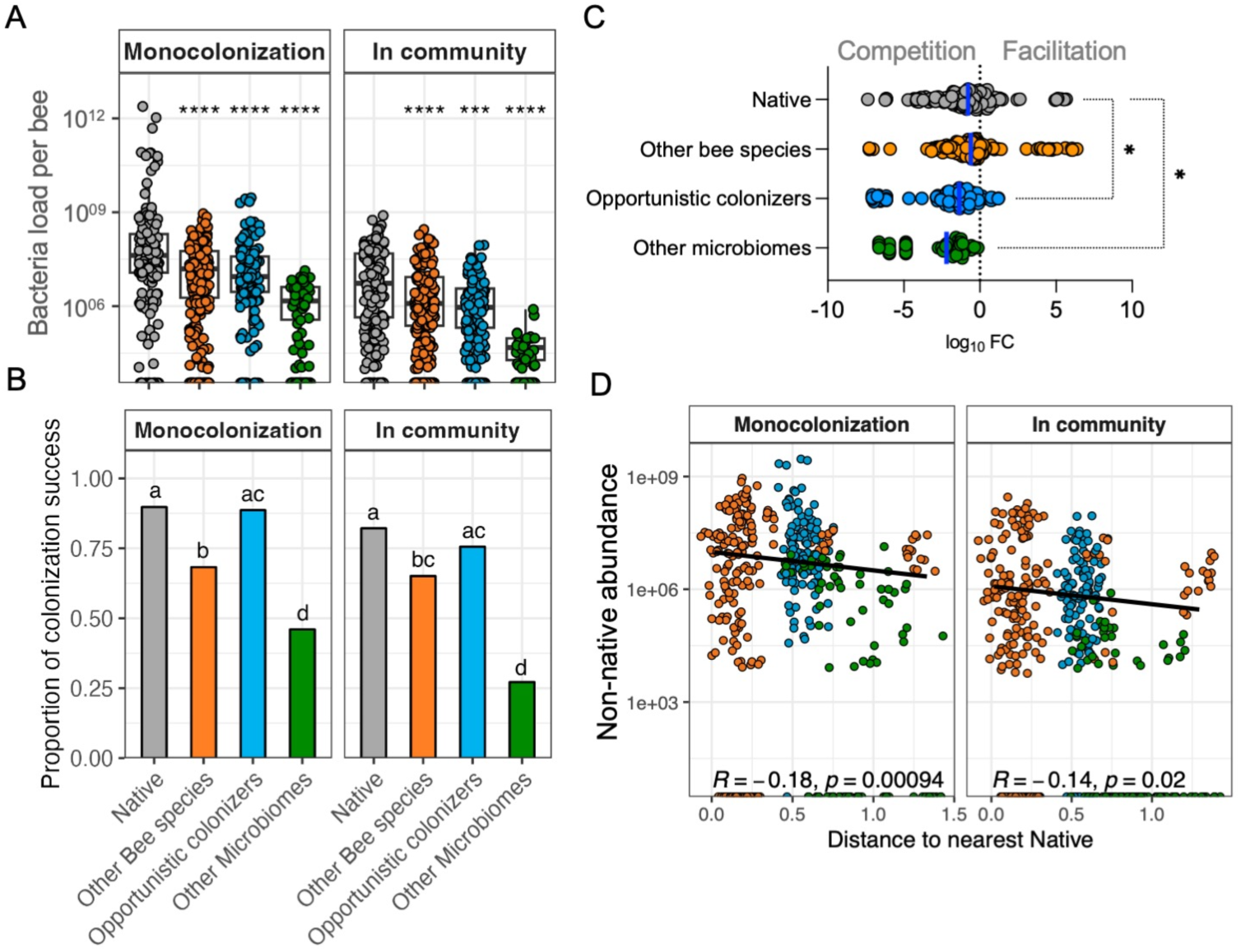
Bacterial categories differ in their absolute loads and overall colonization success. **A.** Abundance of each strain, grouped by isolation source, in the honey bee gut during monocolonization or in the presence of the native community. Pairwise Wilcoxon rank-sum test. P-values were adjusted using FDR method. Asterisks denote pairwise comparisons to the *Native* category within each context (mono or SC). For context, total bacterial loads in microbiota-depleted (MD) and native community–colonized (SC) control bees are shown in Fig. S1. **B.** Proportion of colonization success of each strain, grouped by isolation source, in the honey bee gut during monocolonization or in the presence of the native community. Different lowercase letters indicate significant differences based on Chi-square for equality of proportions and p-value FDR-corrected for multiple comparisons. Community: χ²(3)=99.7, *p*<2.2×10⁻¹⁶; estimates: 0.82 vs 0.65 vs 0.76 vs 0.27; FDR pairwise: native > other-bee, *q*=3.9×10⁻⁴; other microbiomes < all, *q*≤1.2×10⁻¹⁰; native ≈ opportunistic, *q*=0.21. Bars show pooled proportions within each category (Σsuccess / Σtotal; n=12 bees per strain. **C**. Difference in abundance of each strain between monocolonization and in the presence of the native community, grouped by isolation source, and expressed in log10 fold change (log10FC). log10FC < 0 indicates competition (lower abundance in community), whereas log10FC > 0 indicates facilitation (higher abundance in community. Kruskal-Wallis rank sum test, FDR corrected p-value. The mean log10FCFC shows that competition is significant in all categories based on one sample Wilcoxon test, FDR corrected. P value < 0.0001 for all categories. Significance labels as p<=0.0001:****, p<=0.001:***, p<=0.05:*. **D.** Differences in abundance of non-native strains correlate with phylogenetic distance to their nearest native strain. Phylogenetic distances are the horizontal distances in the phylogenetic tree (Figure 1A). R depicts Spearman correlation coefficient. Only non-native strains are shown (native strains are excluded); point colors indicate ecological category.

Given the observed variation in colonization among strains, we tested whether phylogenetic relatedness to native gut symbionts of *Apis mellifera* could account for differences in colonization. Consistent with this hypothesis, we found a significant negative correlation between phylogenetic distance to the closest native strain and colonization abundance among non-native strains for both monocolonization and community context (Figure 2D, Figure S3). Overall, these findings demonstrate that while the ability to colonize the honey bee gut is not limited to native strains, native symbionts generally exhibit higher colonization loads.

### KEGG-based metabolic profiles reveal category-specific rather than universal colonization signals

To investigate whether metabolic capacity underlies differences in colonization between strains, we evaluated the completeness of KEGG functional modules across all bacterial strains. A total of 40 KEGG metabolic pathways were identified, and module completeness was quantified per strain (Figure S4, supplementary File S4). A PCA of KEGG module completeness did not separate strains by colonization performance: strains with high and low gut loads were intermixed (Figure 3A), and the same pattern was observed for colonization efficiency (% colonization; Figure S5). Hierarchical clustering based on KEGG modules grouped several *Native* and *Other bee species* strains together, especially those with well-represented central carbohydrate metabolism, but this clustering did not reflect colonization outcome (Figure S6). We then asked whether specific metabolic functions were linked to colonization performance. A KEGG module-specific mixed-effects scan for colonization abundance and efficiency identified only a small number of modules with significant associations (p < 0.05), and these effects were modest. Positively associated modules were mostly amino-acid-related, with lysine biosynthesis linked to higher abundance and tyrosine, theorine, and leucine pathways linked to higher colonization efficiency (Figure 3B, supplementary file S4). In contrast, several stress- or envelope-related modules showed negative coefficient values in the abundance model. Modules linked to cell-wall remodeling or antibiotic-resistance-type functions (e.g. vancomycin-type D-Ala-D-Lac remodeling) and to siderophore/iron acquisition were associated with modest but significant reductions in bacterial load. This pattern was not observed for colonization efficiency, where the main signals were amino acid and other anabolic pathways with positive effects (Figure 3B, supplementary file S4). Although metabolic completeness did not universally predict colonization capacity across all ecological groups, we observed significant category-specific trends. Specifically, a positive correlation was found between the number of high-completeness KEGG modules and median colonization abundance for strains from *Other microbiomes*, while a negative correlation was observed for *Native* strains (Figure 3C). However, the overall number of KEGG modules with ≥80% completeness did not significantly differ between ecological categories (Figure 3D).

**Figure 3.**
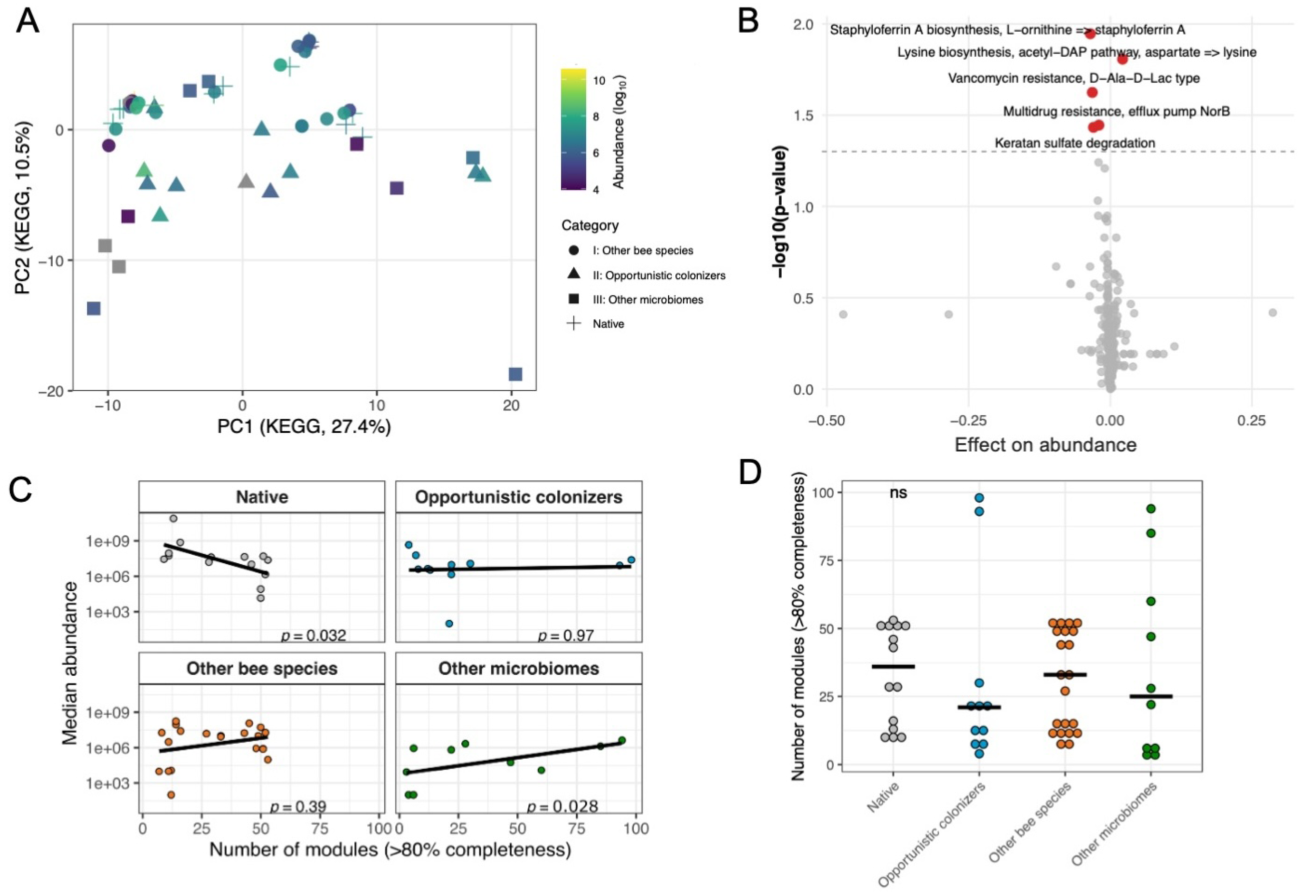
Metabolic capacity of the tested strains across the four isolation categories. **A.** PCA of KEGG module completeness across strains. Principal component analysis was performed on KEGG module completeness profiles. Each point is a strain, positioned by PC1 (27.4%) and PC2 (10.5%), colored by abundance in monocolonization. The shape indicates the category. **B.** Association between KEGG module presence and bacterial abundance. Each point is one KEGG module from the mixed-effects model (model: log_median ∼ module + Category + Colonization_mode + (1 | genus)). The dashed line marks p = 0.05. The model adjusts for category and genus. Points in red are modules with p-value < 0.05. **C.** Relationship between strain abundance and metabolic module content. Median abundance of each strain in monocolonization is plotted against its total number of KEGG modules with ≥80% completeness. Points are coloured by colonization category. A rank-based mixed-effects model was fitted accounting for category and genus: log_median_rank ∼ Number.of.modules_rank + Category + (1 | genus). Isolation category explained part of the variation, whereas there was minimal genus-level variance (intercept for genus was estimated at 0) for n = 56 strains from 25 genera. **D.** Metabolically complete modules by isolation category. Number of KEGG modules ≥80% complete per strain, compared across categories using a mix model (n_mod80 ∼ Category + (1 | genus)).

### Differential activation of host immune effectors by individual gut bacterial strains

To evaluate how different bacterial colonizers influence host immune responses, we mono-associated MD honey bees with 25 of the 56 strains and measured the expression of seven immune-related genes (Apidaecin1, Abaecin, Defensin1, Defensin2, Hymenoptaecin, IRP30, Lysozyme) in the gut using qPCR seven days post-colonization. Strains were selected based on their ability to reach colonization levels >10^6^ bacteria per gut (Figure 1C) and to represent each of the four distinct ecological categories. Expression levels were evaluated relative to MD controls.

Overall, we observed relatively few significant changes in immune gene expression between MD bees and those colonized with bacteria. For instance, only *Apidaecin 1* and *Defensin 2* were consistently upregulated in bees colonized with the full native community compared to MD controls. Likewise, only a limited number of monocolonizations led to detectable changes in gene expression (Figure S7A, Supplementary File S5). AMP and *Lysozyme* expression levels varied considerably across individual bacterial strains, but these differences did not segregate by ecological category; strains from the same group (e.g. native *Gilliamella*, native *Lactobacillus*) often elicited divergent responses (Figure S7A). While certain non-native strains triggered elevated expression of *Apidaecin 1*, *Defensin 2*, and *Lysozyme*, others within the same group did not. Similarly, some native strains also elicited measurable immune responses, while others did not. No consistent expression patterns were observed for *Hymenoptaecin* or *IRP30* across any colonizer group. Exceptions to this general trend included *Defensin 2*, which was more frequently upregulated by native strains, and *Lysozyme*, which showed the opposite trend, being more often induced by non-native strains (Figure S7B). However, overall, immune gene expression responses appeared primarily strain-specific rather than determined by ecological origin.

To further examine whether immune gene expression levels are associated with colonization efficiency, we correlated the median abundance of each bacterial strain in monocolonization with expression of the seven immune genes. No significant correlations were observed (Figure S8), though *Lysozyme* showed a weak negative trend (R=-0.34, p=0.08). Thus, variation in host immune gene expression at seven days post-colonization does not reliably predict bacterial colonization efficiency.

### Sensitivity of native and non-native strains to host AMPs

To better understand the factors influencing colonization in the honey bee gut, we investigated bacterial sensitivity to host-derived AMPs, including Apidaecin1, Abaecin, Defensin1, Defensin2, and Hymenoptaecin, as these immune effectors are known to act selectively on microbes and could differentially inhibit strains depending on their resistance capabilities. To evaluate the potential role of host AMPs in shaping colonization outcomes, a subset of native and non-native strains was screened *in vitro*, and sensitivity scores were derived based on growth inhibition across AMP dilution series, with higher scores reflecting greater susceptibility (Figure 4A, Supplementary File 6). *Gilliamella* strains were among the most AMP-sensitive, with all five tested strains showing susceptibility to at least three AMPs. Similarly, all three *Snodgrassella* strains were sensitive to three or more AMPs, though their overall sensitivity scores were lower than those of *Gilliamella*. Among opportunistic colonizers, *Hafnia* also exhibited high sensitivity, being inhibited by at least three AMPs. Notably, strains of *Apibacter* and *Bifidobacterium* isolated from *Apis cerana* also showed broad AMP susceptibility, each susceptible to at least three antimicrobial peptides. Strains from both the *Native* and *Other bee species* categories exhibited higher AMP sensitivity scores compared to other groups, with strains from the *Other bee species* category showing the highest overall sensitivity (Figure 4B). We next examined whether AMP sensitivity influenced colonization efficiency and colonization success. The median abundance of AMP-sensitive strains was negatively correlated with total sensitivity score in both monocolonization and community contexts (Figure 4C). However, the percentage of colonization success events did not significantly correlate with AMP sensitivity (Figure 4C). Because immune tolerance is likely shaped by host adaptation, we investigated whether bacterial sensitivity to AMPs is related to their phylogenetic relatedness to native strains. We found that total AMP sensitivity scores were significantly negatively correlated with phylogenetic distance to native strains (Figure 4D), indicating that non-native bacteria more closely related to native symbionts tend to be more sensitive to host immune effectors.

**Figure 4.**
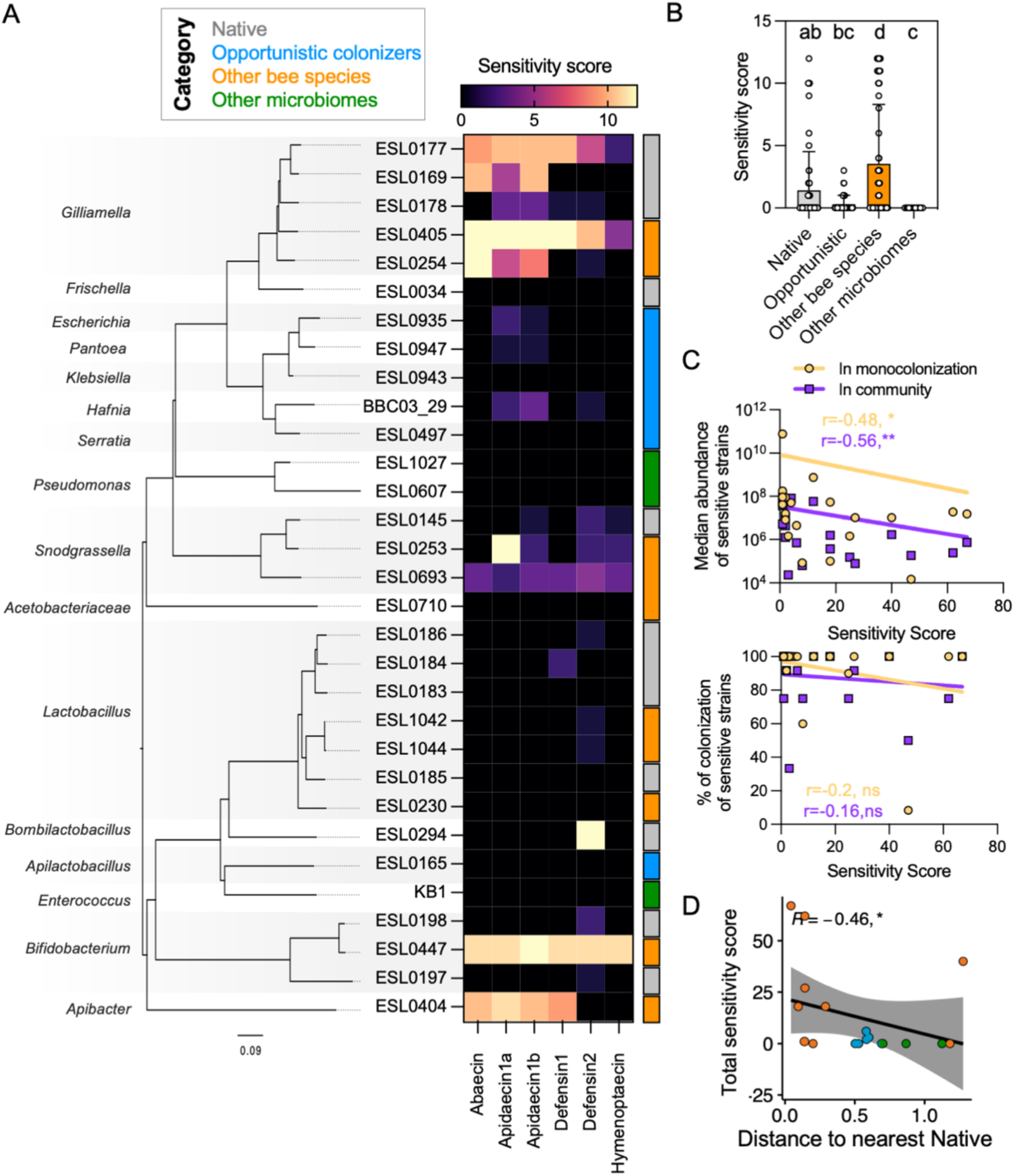
Sensibility of native and non-native strains to host AMPs. **A.** On the left: phylogenetic tree of strains tested for AMP sensitivity inferred by OrthoFinder using the STAG algorithm^51^. On the right: Heatmap of the sensitivity score for a subset of strains in all four categories. Growth curves were performed with 12 dilution series for each peptide that ranged from 100 ug/ml to 0.0487 ug/ml. Experiments were performed in triplicates. Sensitivity score ranged from 0 (not significant growth defects) to 12 (significant growth defects in the lowest concentration). See Material and Methods for details. Supplementary File S6, Figures S9-S14. **B**. Sensitivity scores per isolation category. Different lowercase letters indicate significant differences based on Kruskal-Wallis rank test with p-values FDR-corrected for multiple comparisons. **C.** Upper panel shows the correlation between the median abundance and the total sensitivity score of each strain in monocolonization (yellow) and in the presence of the native community (purple). Lower panel shows the same but for the percent of colonization success. **D.** The total sensitive score was calculated as the sum of the individual sensitivity scores for each tested peptide. R depicts the Spearman correlation coefficient values. Significant labels: p<=0.01:**, p<=0.05:*. Data points are colored by category according to color-code in panel A.

## DISCUSSION

In this study, we examined how host filtering mechanisms contribute to the specificity of the gut microbiota in social bees. We tested an ecological and phylogenetically diverse set of bacterial strains, and found that, on average, native strains achieved higher bacterial loads and colonized a greater proportion of hosts in both mono-association and in the presence of the native microbiota. This is consistent with some previous work on host specificity, which report pronounced home-site advantages among bee gut core symbionts (e.g., *Snodgrassella*^30^, *Gilliamella*^39^, *Lactobacillus*^32^). In mouse models, native *Bacteroidetes* and *Firmicutes* strains similarly show home-site advantages, outcompeting non-native counterparts even in hosts lacking an adaptive immune system^10^. Together, these findings suggest that intrinsic host factors preferentially support resident lineages. Despite this overall trend, we also observed substantial variation in colonization success even among closely related native strains, highlighting the importance of testing host specificity across a diverse set of strains.

Moreover, some bacteria from environments outside the animal gut were able to colonize honey bees to a certain extent, and even in the presence of the native microbiota, most of the non-native strains still achieved some level of colonization. However, in most cases, their colonization success was significantly reduced. This suggests that host filtering is not entirely strict, and also that non-native strains are not completely outcompeted by native strains when introduced at sufficiently high numbers. Related findings in other host systems show that environmentally derived or heterologous host-associated bacteria can colonize, and in some cases even dominate, gnotobiotic or germ-free hosts^14,53^.

Host-specific colonization of bee gut symbionts likely reflects interactions among ecological, bacterial, and host factors. In honey bees, early social acquisition favors nestmate-associated bacteria, and fecal transplants can transfer a gut-like microbiota to microbiota-depleted bees^35^. Consistently, reduced colonization of both native and non-native strains in the presence of the native community (Figure 2C) indicates strong competition for gut establishment. Resource limitation, niche preemption, and antagonism may therefore allow resident, socially transmitted strains to exclude environmental or non-native bacteria, reinforcing host specificity, as seen in other systems^19,54,55^.

Because the negative relationship between colonization load and distance to the closest native strain was also observed in monocolonization (Fig. 2D), it cannot be explained solely by competition with resident close relatives. Instead, phylogenetic proximity likely reflects host compatibility, that is, conserved traits enabling persistence and growth under host-derived constraints, consistent with the association between AMP sensitivity and colonization load (Fig. 4C-D). Together with the higher colonization success of native strains, this indicates that bacterial traits and host filtering are key determinants of host specificity. One possible factor is metabolic adaptation to diet- or host-derived nutrients in the honey bee gut. Although utilization of pollen-derived carbohydrates is an important trait of bee gut symbionts^56,57^, non-native strains with similar functional profiles did not necessarily colonize better (Figure 3, Figures S5-S6). Indeed, only a small subset of amino acid-related modules was positively, and only modestly, associated with colonization. In addition, the negative association between KEGG module richness and colonization observed among native strains (Fig. 3C) may partly be driven by the representation of specific bacterial lineages within this group, particularly high-colonizing Lactobacillus/Bombilactobacillus strains that show comparatively lower KEGG module richness (Fig. S5), consistent with genome streamlining reported for these host-specialized bee gut symbionts^32,58^. An important limitation of this analysis is that KEGG annotation provides only a coarse-grained functional profile of a bacterium, with many functions either omitted or described only at a general level. Module completeness may be underestimated in fragmented assemblies. Differences in gene regulation and in the expression of metabolic pathways may further modulate colonization success, irrespective of the mere presence of these functions within the genome.

Another important factor that may contribute to host filtering is the host’s immune system. In our study, host immune gene expression in response to gut colonization appeared to be primarily strain-specific rather than clearly associated with a particular ecological category of the tested strains (Figure S7A). Even among native taxa, closely related strains differed in their capacity to stimulate bee immunity, mirroring previous reports that some native *Lactobacillus* strains can markedly upregulate AMP genes, whereas others do not^59^. Apidaecin was often strongly induced by colonization with the full native community or even a single native strain, while other AMPs (Abaecin, Defensin) changed little^38^. A niche-matched contrast underscores this strain specificity: although *Frischella perrara* and *Snodgrassella alvi* both colonize the ileum, *F. perrara* triggers substantially stronger AMP and immune-pathway response than *S. alvi*^60^. Such differences may reflect not only bacterial traits but also the specific gut region sampled, such as midgut, pylorus, or the full hindgut, each of which exhibits distinct patterns of bacterial colonization and immune activity^36,61^. Overall, we observed relatively few differences in immune gene expression between colonized and MD bees seven days post-inoculation, except for *Abaecin* and *Defensin 2* (Figure S7A). This is consistent with findings from Kwong et al^38^., who similarly reported limited immune gene induction five days post-colonization. It is possible that by these relatively late time points following bacterial colonization, the immune response to bacterial exposure had largely subsided. Indeed, several previous studies which assessed earlier stages of colonization, such as 12 h or 24 h post-inoculation, reported the expression of additional immune-related genes through both the immune deficiency (IMD) and Toll signaling pathways^37,39,59^ that may be involved in the recognition of native versus non-native bacterial strains. The fact that we did not observe any significant correlation between immune gene expression and bacterial abundance (Figure S8) is consistent with the idea that immune responses to bacterial colonization are transient. This temporal dynamic is not unique to honey bees. In other well-studied host-microbe systems, immune and developmental gene expression changes often peak early during microbial establishment and diminish over time. For example, in the *Drosophila*-*Lactobacillus* model and the *Euprymna scolopes*-*Vibrio fischeri* squid symbiosis, aposymbiotic hosts exhibit delayed expression changes compared to symbiotic individuals, particularly during early developmental windows critical for symbiont integration^62–65^. Bacterial signals often modulate the timing rather than the occurrence of host developmental processes: the same changes can happen in symbiotic and germ-free hosts, but they start earlier when microbes are present (e.g. onset of foraging in honey bees, apoptotic arms in squid light organs)^7,66,67^. These comparisons highlight the importance of temporal resolution when evaluating host responses and suggest that early time points may be more informative for capturing the acute immune signaling associated with microbial recognition and integration.

Bacteria can differ not only in their ability to trigger host immune responses but also in their capacity to withstand them, which can influence their colonization success. Our results on AMP sensitivity revealed a high degree of variation in immune tolerance among the tested strains, similar to the variation observed in immune responses, although AMP sensitivity showed a clearer association with the strains’ ecological categories. Bee-associated bacteria generally exhibiting greater susceptibility than bacteria isolated from other environments (Figure 4A, 4B), and non-native strains were overall more sensitive than native strains of the core microbiota (Figure 4B). This is consistent with earlier work showing genus-level differences in AMP sensitivity^38,68^. We found *Snodgrassella* tend to be more resistant to apidaecin 1b than *Gilliamella*, and that Gram-positive core members such as *Lactobacillus* and *Bifidobacterium* show broad resistance to apidaecin and hymenoptaecin. Interestingly, we found a negative correlation between overall sensitivity to the tested AMPs and bacterial load, both in monocolonization and in the community context (Figure 4C). This observation corroborates that AMPs may play an important role in regulating symbiont abundance and maintaining microbial homeostasis. Interestingly, *Gilliamella* strains were among the most sensitive strains, followed by those of the genus *Snodgrassella*. While this might seem paradoxical given their status as successful colonizers, it aligns with previous reports showing that Gram-negative symbionts, particularly *Gilliamella*, are more vulnerable to apidaecins^38,69^. This apparent sensitivity may not reflect a disadvantage, but rather a finely balanced immune-microbe interaction where AMPs act as regulators that maintain microbial homeostasis rather than eliminators of symbionts^70–72^, complementing ROS-mediated mechanisms that exclude maladapted strains^11,39^. Indeed, while AMP sensitivity correlated with bacterial abundance, it did not significantly predict colonization success or failure (Figure 4C, lower panel). Thus, AMPs seem to act in a graded, context-dependent manner that controls symbiont load rather than strictly permitting or blocking colonization, a mode of action also reported in other insects where AMPs confine resident symbionts to specific tissues (e.g. coleoptericin-A in *Sitophilus*^73^).

Finally, we observed a significant inverse relationship between AMP sensitivity and phylogenetic distance from native bee strains (Figure 4D). Strains more closely related to core symbionts of *A. mellifera* were more sensitive to host AMPs. Rather than selecting for broad resistance, the immune system may have evolved to support stable interactions with beneficial microbes by modulating their growth. Immune effectors can mediate not just defense, but the spatial and population-level dynamics of beneficial symbionts, a principle that appears conserved across diverse host taxa^74^.

It is important to recognize that AMP resistance is not the sole determinant of microbial fitness within the host gut. Some native strains that display sensitivity to honey bee AMPs may still thrive due to alternative or compensatory mechanisms. For instance, co-colonization or cross-feeding interactions may buffer AMP-sensitive strains against immune pressure, enabling persistence in the gut despite their inherent susceptibility. AMP-sensitive strains may persist when shielded or supported by compatible community members. This phenomenon has parallels in other systems. In *D. melanogaster*, AMP-sensitive strains are often outcompeted unless co-colonized with more immune-tolerant bacteria that can modulate host immunity or provide spatial refuge^75,76^. Similarly, in mammals, microbial persistence under immune pressure is often mediated by cooperative traits such as biofilm formation, cross-feeding, or immune evasion mechanisms^77,78^. Moreover, the host immune landscape is itself complex and dynamic. Natural polymorphisms in AMP genes among honey bee populations could lead to variability in the selective pressures experienced by microbes in different individuals or colonies.

Understanding these layers of complexity will be important for unraveling the full spectrum of host-microbe compatibility and microbial persistence mechanisms in the gut ecosystem. While we focused here on single peptide effects, natural host environments feature mixtures of AMPs that may act synergistically or antagonistically, further shaping community dynamics in complex and non-linear ways^79^. Moreover, other arms of the host immune system such as the DUOX-reactive oxygen species pathway, are activated by gut bacteria in insects and can limit microbial proliferation^39,80,81^. In addition, non-immune system factors, such as access to specific gut niches (pylorus vs ileum vs rectum), attachment sites and competition within these microenvironments, motility, chemotaxis, are likely to play to play a relevant role in host filtering, which we have not explored in this study.

Taken together, our findings support a model in which colonization success results from multifaceted processes with interplay between host immune filtering and microbial ecological dynamics. Early in adult life, socially transmitted core symbionts gain priority access to gut niches and establish a resident community that acts as an ecological filter, suppressing most incoming strains through competitive exclusion, with only rare facilitation. At the same time, host antimicrobial peptides impose a second selective pressure that limits bacterial proliferation in a taxon- and strain-dependent manner, shaping persistence and abundance rather than uniformly blocking establishment. Together, priority effects, microbial interactions, and host immune constraints help explain how honey bees maintain a stable, host-specific adult gut microbiota despite continual exposure to environmental and non-native bacteria.

## ACKNOWLEDGMENTS

We thank the Engel lab for valuable discussions and support throughout this work. In particular, we are grateful to Garance Sarton-Lohéac for generously sharing her genome assembly pipeline. We thank Vincent Sommerville for assistance with 3D-printing the bee cages. We also thank Konstantin Schmidt, Pascale Vonaesch, Christoph Keel, Yolanda Schaerli and Carlos Gustavo Nunes da Silva for providing bacterial strains used in this study. We thank the FBM décanat for the support provided during the parental leave of S.M.G. This work was funded by an SNSF Consolidator grant (’GLOBEE,’ grant number 213860), and the NCCR Microbiomes, a National Centre of Competence in Research (grant number 180575 and 225148), all funded by the Swiss National Science Foundation.

## AUTHOR CONTRIBUTIONS

SMG: Conceptualization, Methodology, Investigation, Formal analysis, Visualization, Writing-original draft, Writing-review & editing. AuP: Investigation, AC: Investigation, Conceptualization, Writing-review & editing. AP: Formal analysis. AS: Investigation, MG: Investigation, LK: Investigation, Writing-review & editing, FM: Conceptualization, Writing-review & editing PE: Conceptualization, Funding acquisition, Resources, Writing-original draft, Writing-review & editing.

## DATA AVAILABILITY

All sequencing data and assembled genomes have been uploaded in the NCBI Sequence read archive repository under the accession number PRJNA1364849. All analysis code is available on GitHub: https://github.com/mogusi8/Host-and-microbial-factors-influence-gut-bacteria-in-honey-bees/tree/main. All other relevant data generated during this study are included in this article and its supplementary files.

## SUPPLEMENTARY MATERIAL

**Figure S1:**
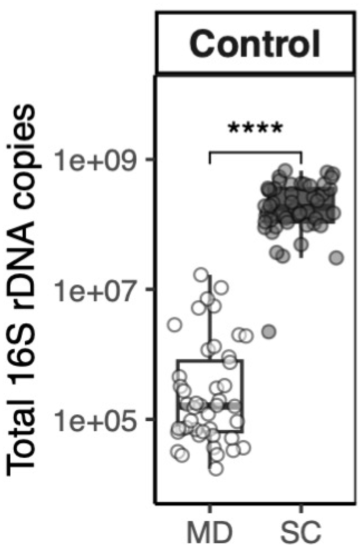
Total bacterial load in control groups (MD and SC) seven days post-inoculation. Total 16S rRNA gene copies per gut were quantified by qPCR using universal bacterial primers, in microbiota depleted (MD) bees and native synthetic community (SC) colonized bees. Mann–Whitney test, n=48. Significant label: p<=0.0001:****.

**Figure S2.**
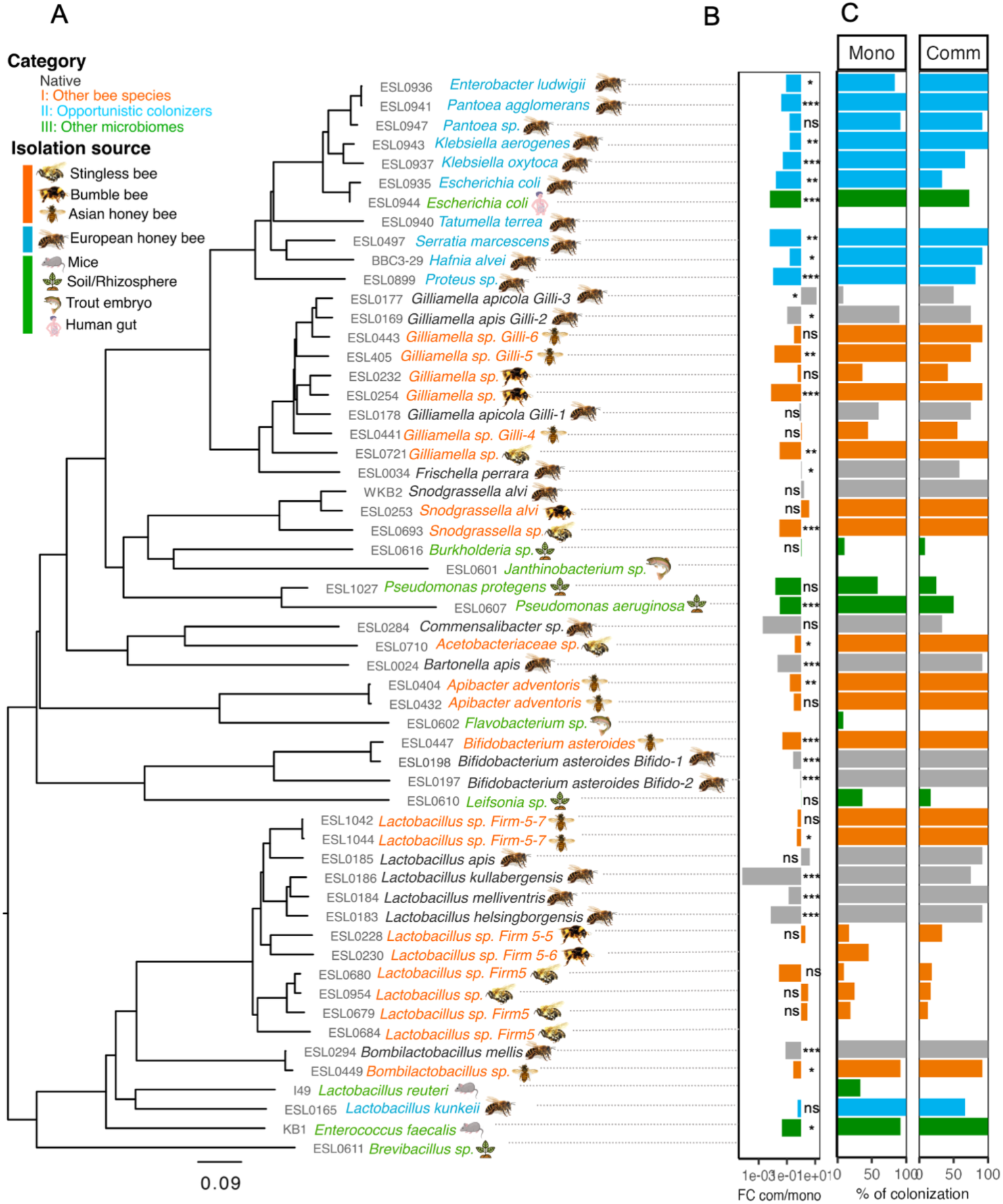
Bacterial colonization success of the honey bee gut, seven days post-inoculation, in monocolonization or in the presence of the native synthetic community (SC). **A** Phylogenetic tree of the 56 bacterial species and strains tested for colonization of the honey bee gut. The tree was inferred with OrthoFinder using the STAG algorithm^51^. **B.** Difference in abundance (expressed in log10 fold change) of each strain between monocolonization and in the presence of the native community. Mono and Comm data were obtained from independent bees; the fold-change shown connects the same strain measured under the two conditions. Wilcoxon rank-sum test. The p-values were adjusted using FDR method. Significance labels shown as p<=0.05:*, p=<=0.01:**, p<=0.001***, ns: not significant. **C.** Percent of colonization success for each strain in monocolonization (Mono) and in the presence of the native community (Comm) based on qPCR absolute abundance calculated as genome equivalents. The gut of a bee was considered colonized when the abundance was higher than in the initial inoculum (see material and methods). All colonization experiments were conducted with n=12 bees, obtained from three independent experimental replicates conducted with bees from different bee hives.

**Figure S3:**
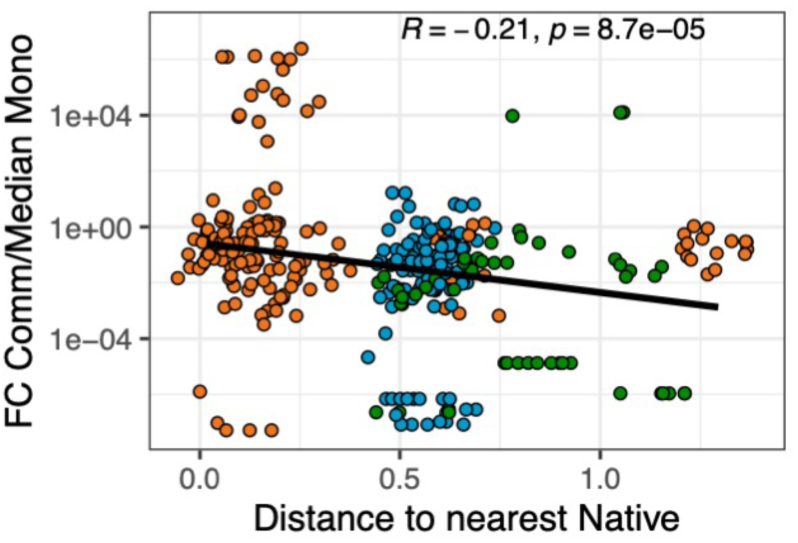
Fold change (FC) in the load of non-native strains in monocolonization relative to colonization in community, plotted against the phylogenetic distance to their nearest native strain. Phylogenetic distances are the horizontal distances in the phylogenetic tree (Figure 1A). R depicts Spearman correlation coefficient. Point colors indicate ecological category (as defined in Fig. 1A).

**Figure S4:**
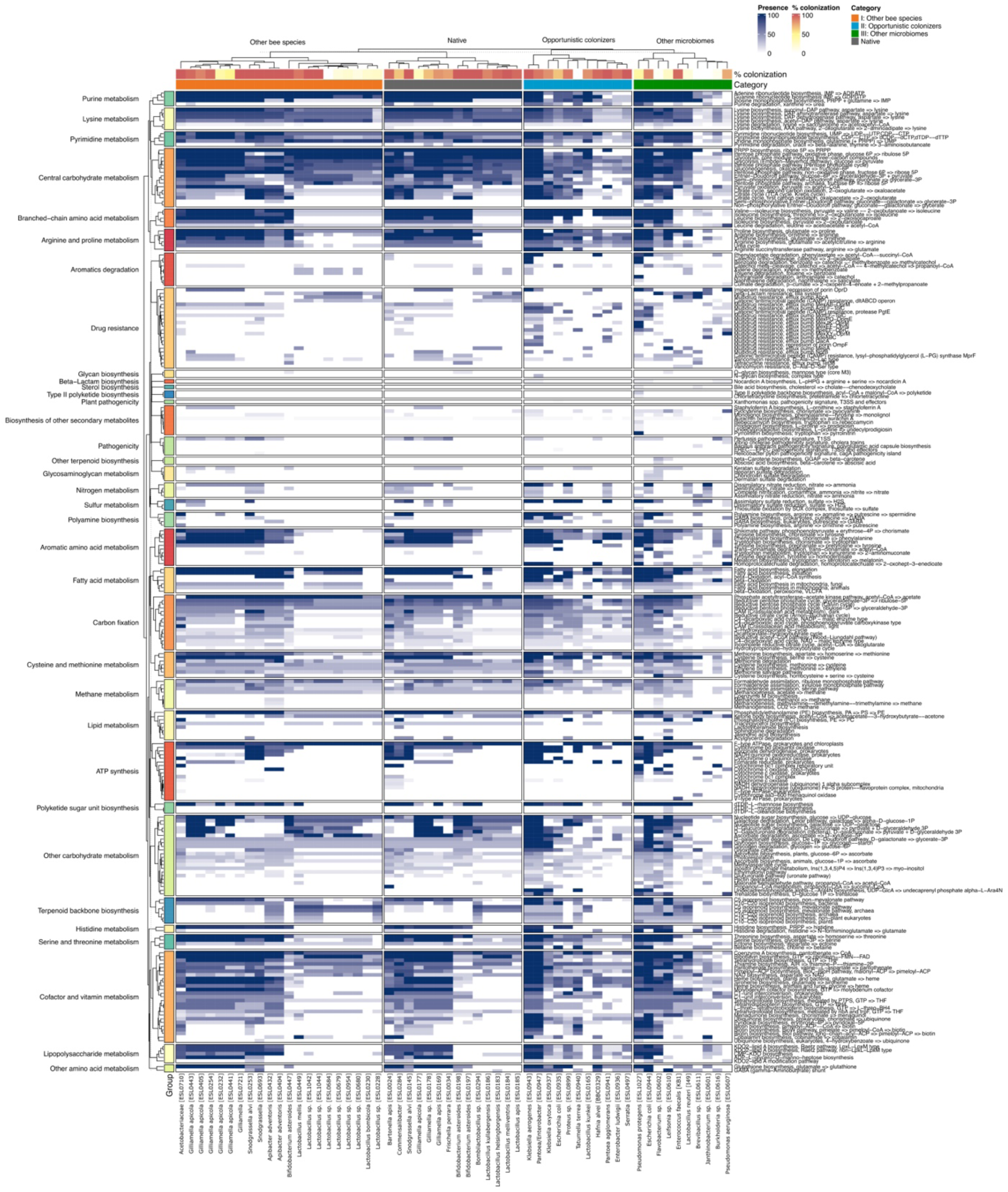
The presence and completeness of KEGG modules in the genomes of individual strains. Heatmap of percent of completeness for each strain. Supplementary file S4.

**Figure S5:**
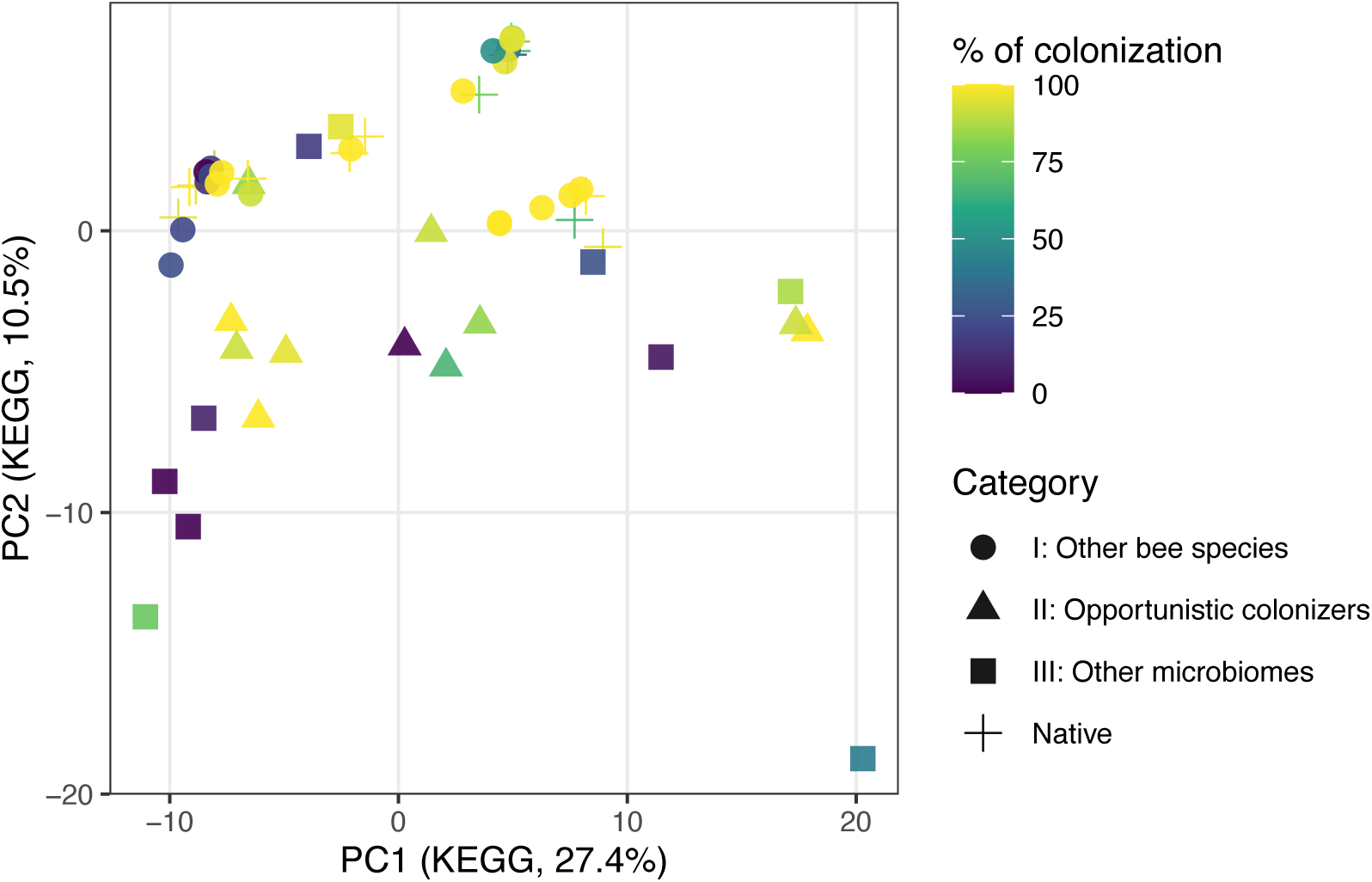
PCA of KEGG module completeness across strains. Principal component analysis was performed on KEGG module completeness profiles. Each point is a strain, coloured by its colonization efficiency in monocolonization. The shape indicates the isolation category.

**Figure S6:**
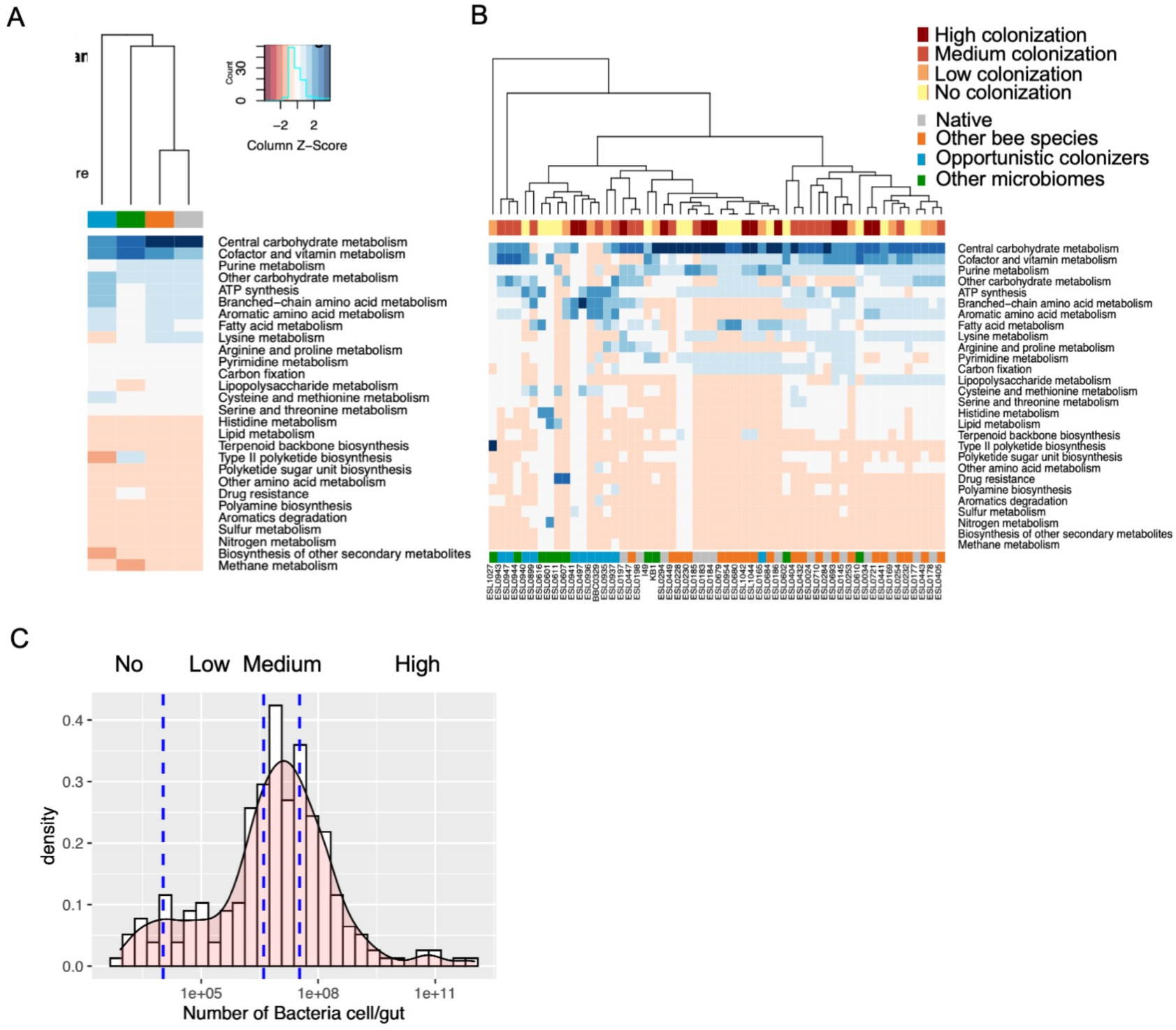
**A.** The presence and completeness of KEGG modules in the genomes of individual strains categorized by isolation source. Heatmap of % completeness for each strain in each category. Grey: *Native*; orange: *Other bee species*; blue: *Opportunistic colonizers*; green: *Other microbiomes*. **B.** Heatmap of the number of modules that are at least 80% complete for each pathway group. Top label shows bacterial loads categorized in for groups based on the abundance distribution in monocolonization (See materials and methods). **C**. Distribution of bacterial loads in honey bee guts during monocolonization. Strains are categorized based on colonization levels, defined by load percentiles: *No colonization* (below the 25th percentile), *Low colonization* (25th-50th percentile), *Medium colonization* (50th-75th percentile), and *High colonization* (above the 75th percentile).

**Figure S7:**
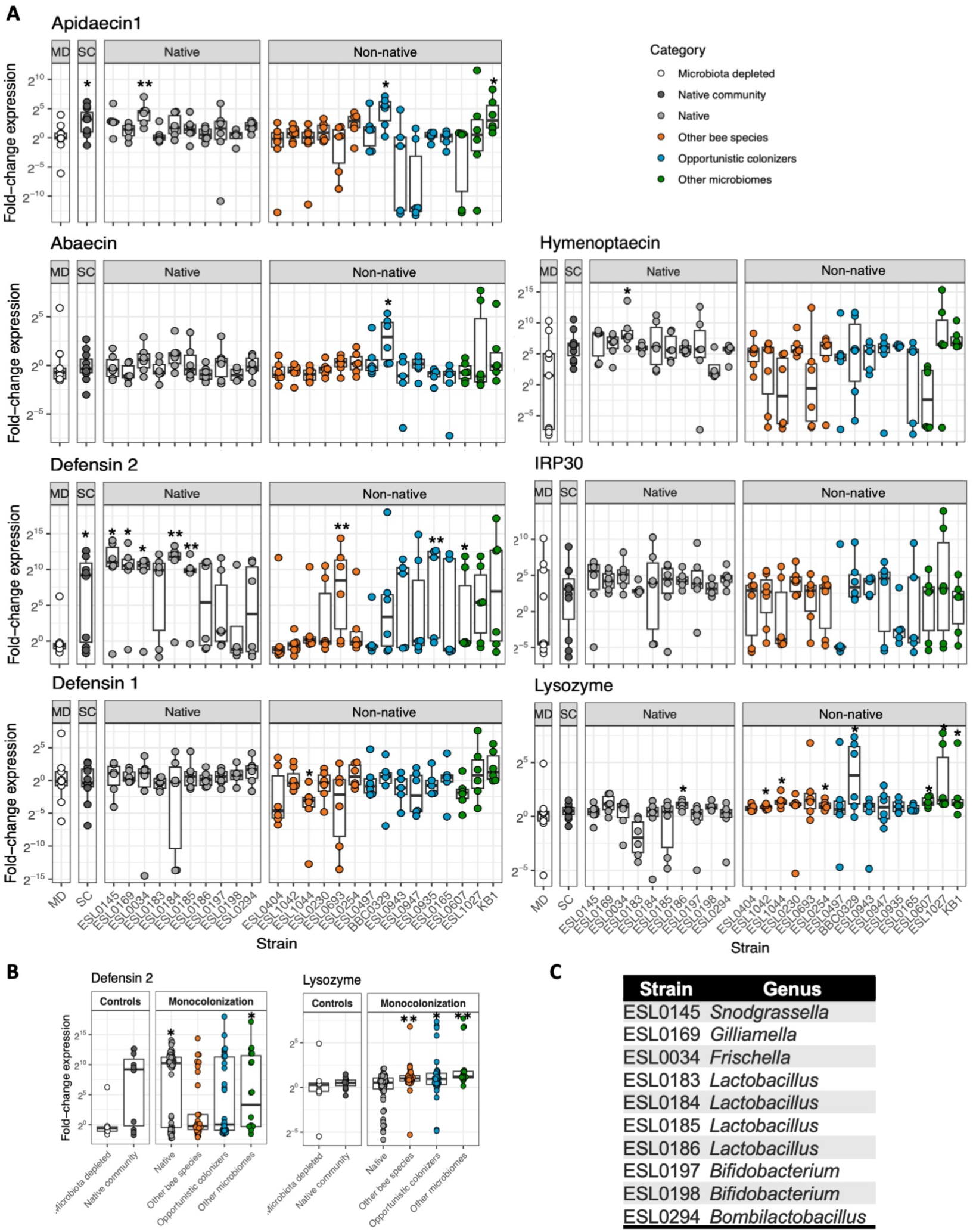
Host antimicrobial peptide responses to colonization by individual bacterial strains. **A** Fold-change in expression relative to MD of six antimicrobial peptide (AMP) genes, *Apidaecin1*, *Abaecin*, *Defensin1*, *Defensin2*, *Hymenoptaecin*, *IRP30*, and *Lysozyme,* in the honey bee gut during monocolonization with the indicated bacterial strains. AMP expression was measured relative to microbiota-depleted (MD) bees. Bees colonized with the synthetic native community (SC) are shown as reference controls. Strains are grouped by category and host origin, with separate panels for native and non-native strains. Data are shown as individual points with boxplots indicating the median and interquartile range. Asterisks indicate strains that induced a significant change in expression relative to the MD control (*p* < 0.05, Wilcoxon rank-sum test, FDR-corrected, n=6). Supplementary file S5. **B,** Summary of *Defensin2* and *Lysozyme* expression levels across different strain categories in control (MD and SC) or monocolonization conditions. Asterisks indicate strains that induced a significant change in expression relative to the MD control (*p* < 0.05, Wilcoxon rank-sum test, FDR-corrected). No significant differences in expression were observed among categories for *Apidaecin1*, *Abaecin*, *Defensin1*, *Hymenoptaecin*, or *IRP30*. **C,** Genus corresponding to each Strain ID for Figure S8.

**Figure S8:**
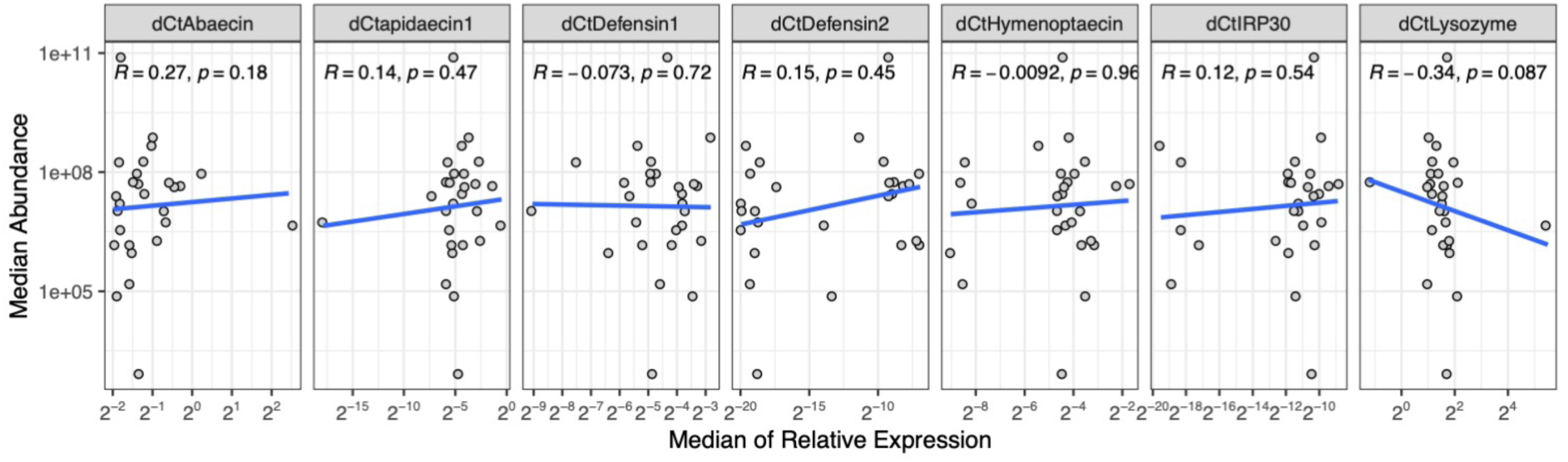
Antimicrobial peptide expression at seven days post-colonization does not predict bacterial colonization efficiency. Correlation between antimicrobial peptide (AMP) expression and bacterial colonization abundance in the honey bee gut. Scatter plots show the relationship between the median relative expression of seven AMP genes (*Abaecin*, *Apidaecin1*, *Defensin1*, *Defensin2*, *Hymenoptaecin*, *IRP30*, and *Lysozyme*) and the median abundance of bacterial strains across samples. Each point represents a bacterial strain. Blue lines indicate linear regression fits. Spearman correlation coefficients (*R*) and associated *p*-values are shown for each gene. Regression lines are shown for visualization only; statistical support for associations is assessed using the Spearman correlation coefficients and p-values reported in each panel.

**Figure S9.**
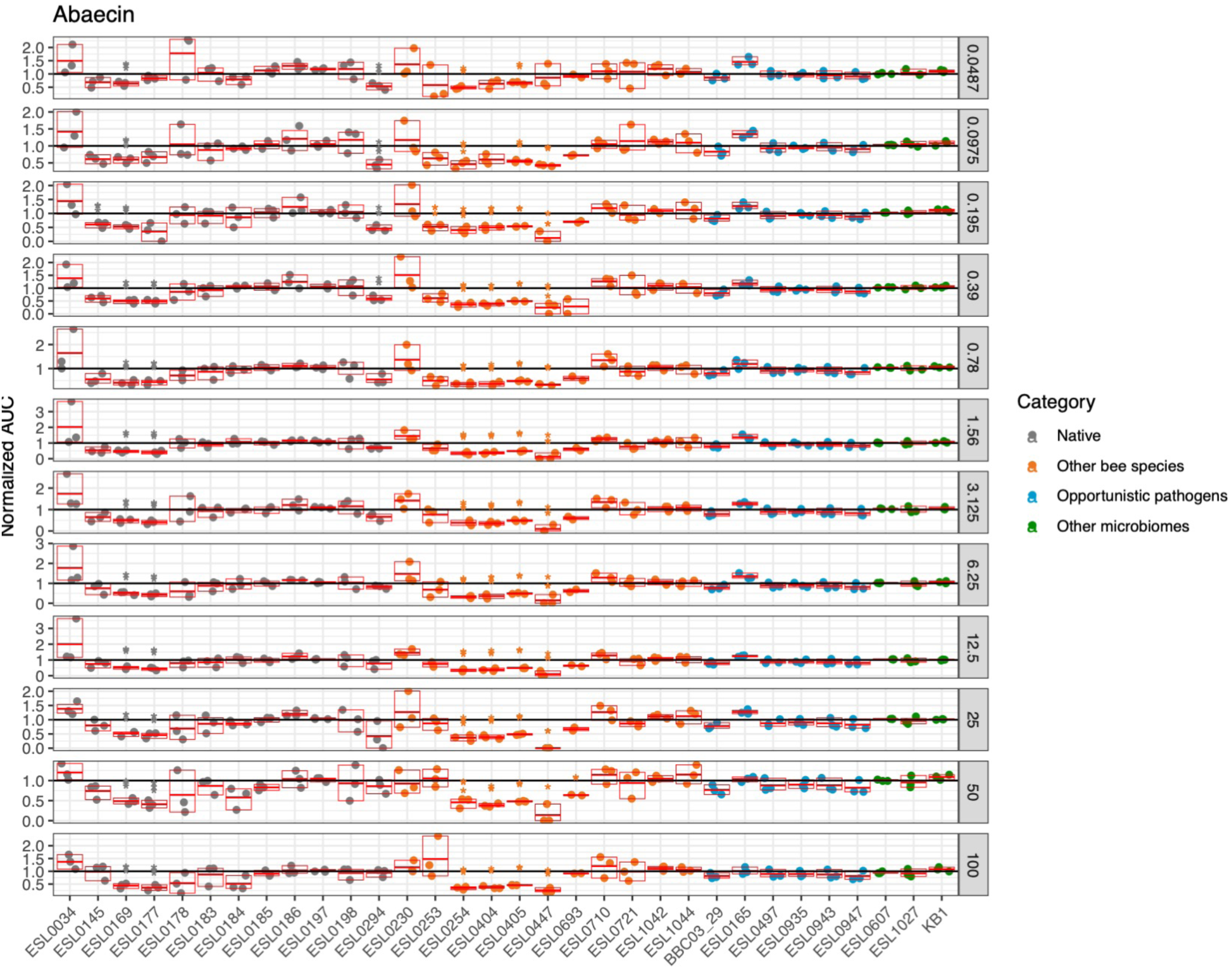
Area under the curve (AUC) of bacterial growth, normalized to the untreated control and presented on a logarithmic scale. The right panel shows the AUC values at a representative concentration of Abaecin. Each condition was tested in triplicate (n = 3). Statistical significance was assessed using a one-sample Wilcoxon test against the null hypothesis of no growth inhibition (normalized value = 1), with p-values adjusted for multiple comparisons using the false discovery rate (FDR) correction. Significance levels are indicated as follows: ****p ≤ 0.0001, ***p ≤ 0.001, **p ≤ 0.01, *p ≤ 0.05.

**Figure S10.**
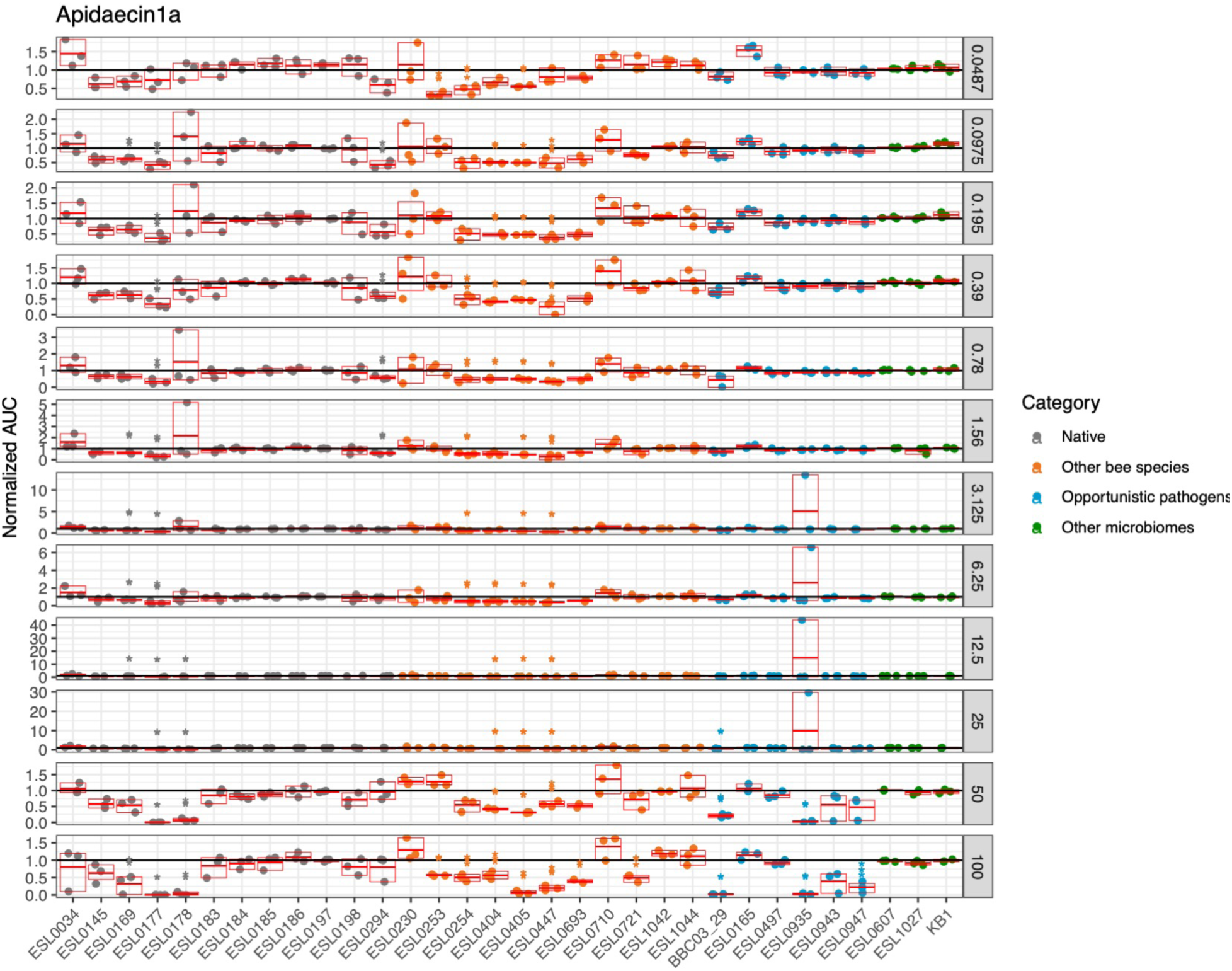
Area under the curve (AUC) of bacterial growth, normalized to the untreated control and presented on a logarithmic scale. The right panel shows the AUC values at a representative concentration of Apidaecin 1a. Each condition was tested in triplicate (n = 3). Statistical significance was assessed using a one-sample Wilcoxon test against the null hypothesis of no growth inhibition (normalized value = 1), with p-values adjusted for multiple comparisons using the false discovery rate (FDR) correction. Significance levels are indicated as follows: ****p ≤ 0.0001, ***p ≤ 0.001, **p ≤ 0.01, *p ≤ 0.05.

**Figure S11.**
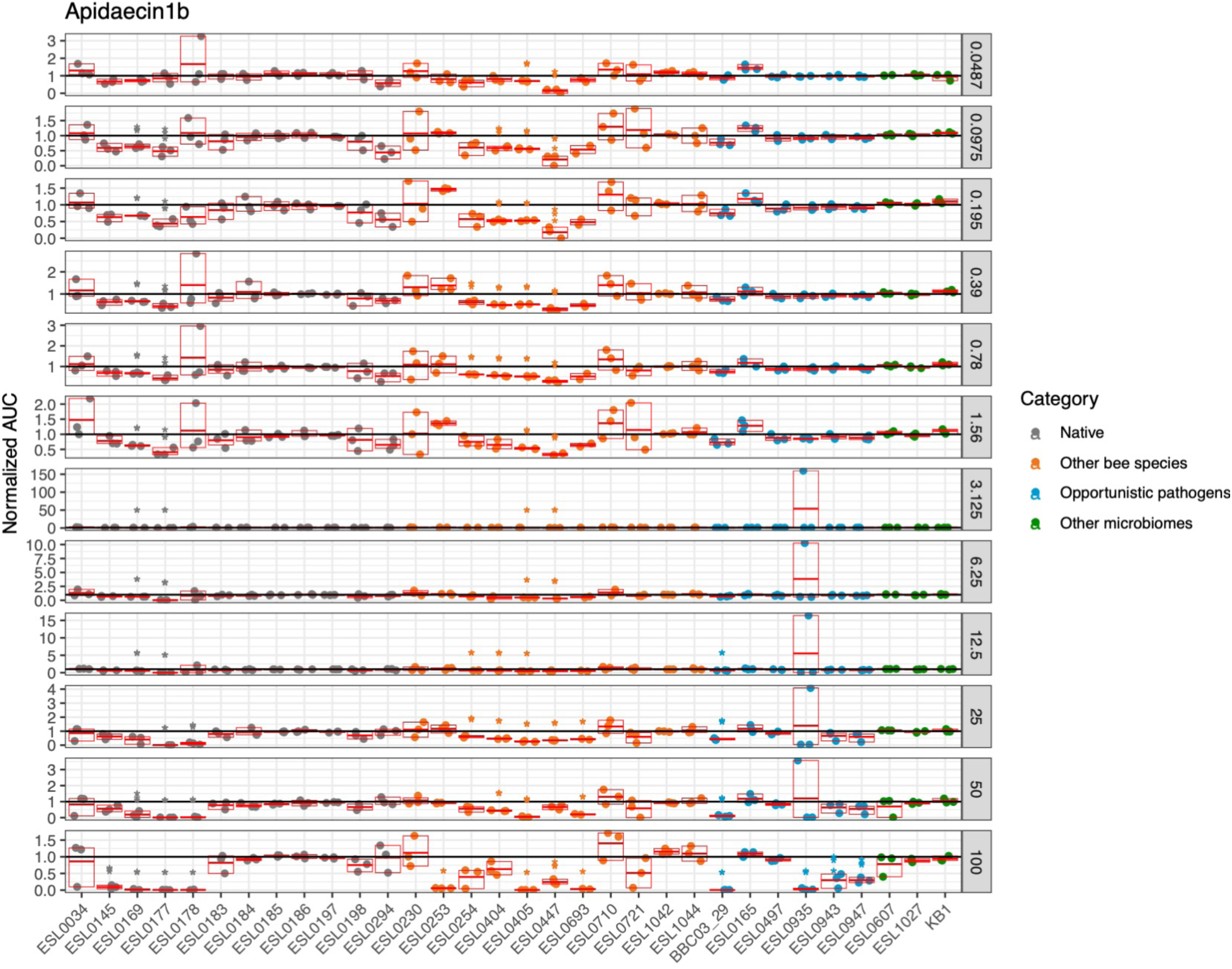
Area under the curve (AUC) of bacterial growth, normalized to the untreated control and presented on a logarithmic scale. The right panel shows the AUC values at a representative concentration of Apidaecin 1b. Each condition was tested in triplicate (n = 3). Statistical significance was assessed using a one-sample Wilcoxon test against the null hypothesis of no growth inhibition (normalized value = 1), with p-values adjusted for multiple comparisons using the false discovery rate (FDR) correction. Significance levels are indicated as follows: ****p ≤ 0.0001, ***p ≤ 0.001, **p ≤ 0.01, *p ≤ 0.05.

**Figure S12.**
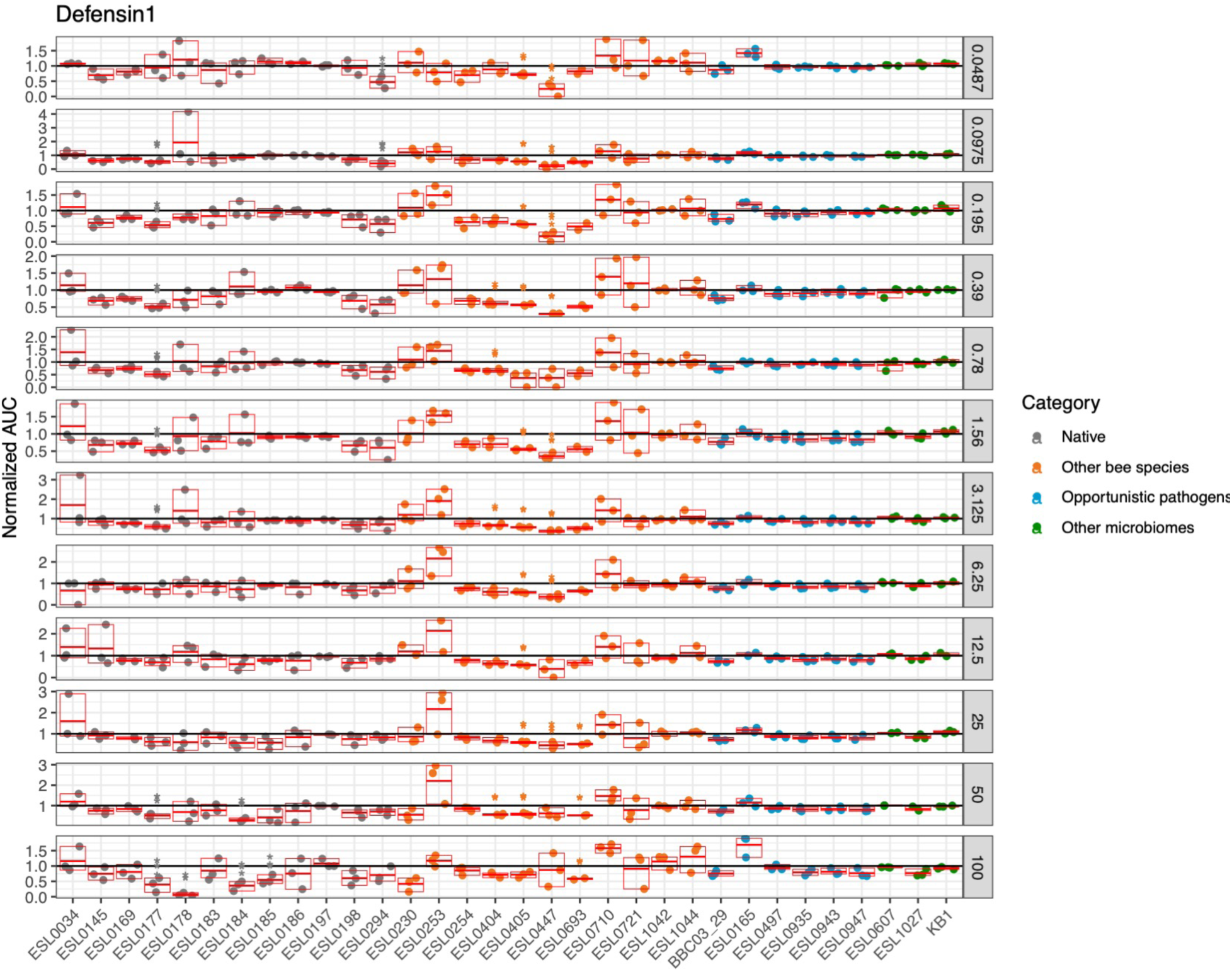
Area under the curve (AUC) of bacterial growth, normalized to the untreated control and presented on a logarithmic scale. The right panel shows the AUC values at a representative concentration of Defensin 1. Each condition was tested in triplicate (n = 3). Statistical significance was assessed using a one-sample Wilcoxon test against the null hypothesis of no growth inhibition (normalized value = 1), with p-values adjusted for multiple comparisons using the false discovery rate (FDR) correction. Significance levels are indicated as follows: ****p ≤ 0.0001, ***p ≤ 0.001, **p ≤ 0.01, *p ≤ 0.05.

**Figure S13.**
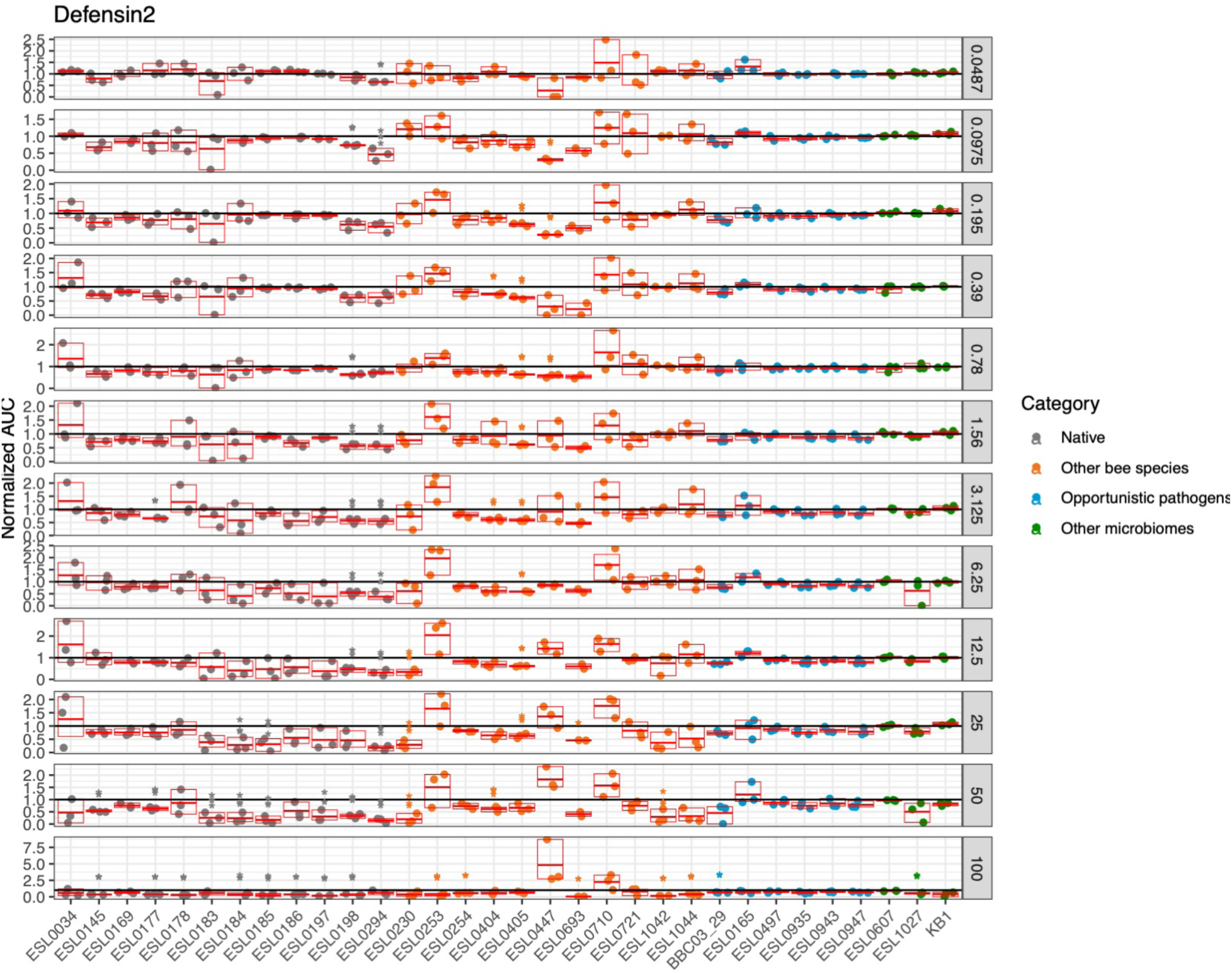
Area under the curve (AUC) of bacterial growth, normalized to the untreated control and presented on a logarithmic scale. The right panel shows the AUC values at a representative concentration of Defensin 2. Each condition was tested in triplicate (n = 3). Statistical significance was assessed using a one-sample Wilcoxon test against the null hypothesis of no growth inhibition (normalized value = 1), with p-values adjusted for multiple comparisons using the false discovery rate (FDR) correction. Significance levels are indicated as follows: ****p ≤ 0.0001, ***p ≤ 0.001, **p ≤ 0.01, *p ≤ 0.05.

**Figure S14.**
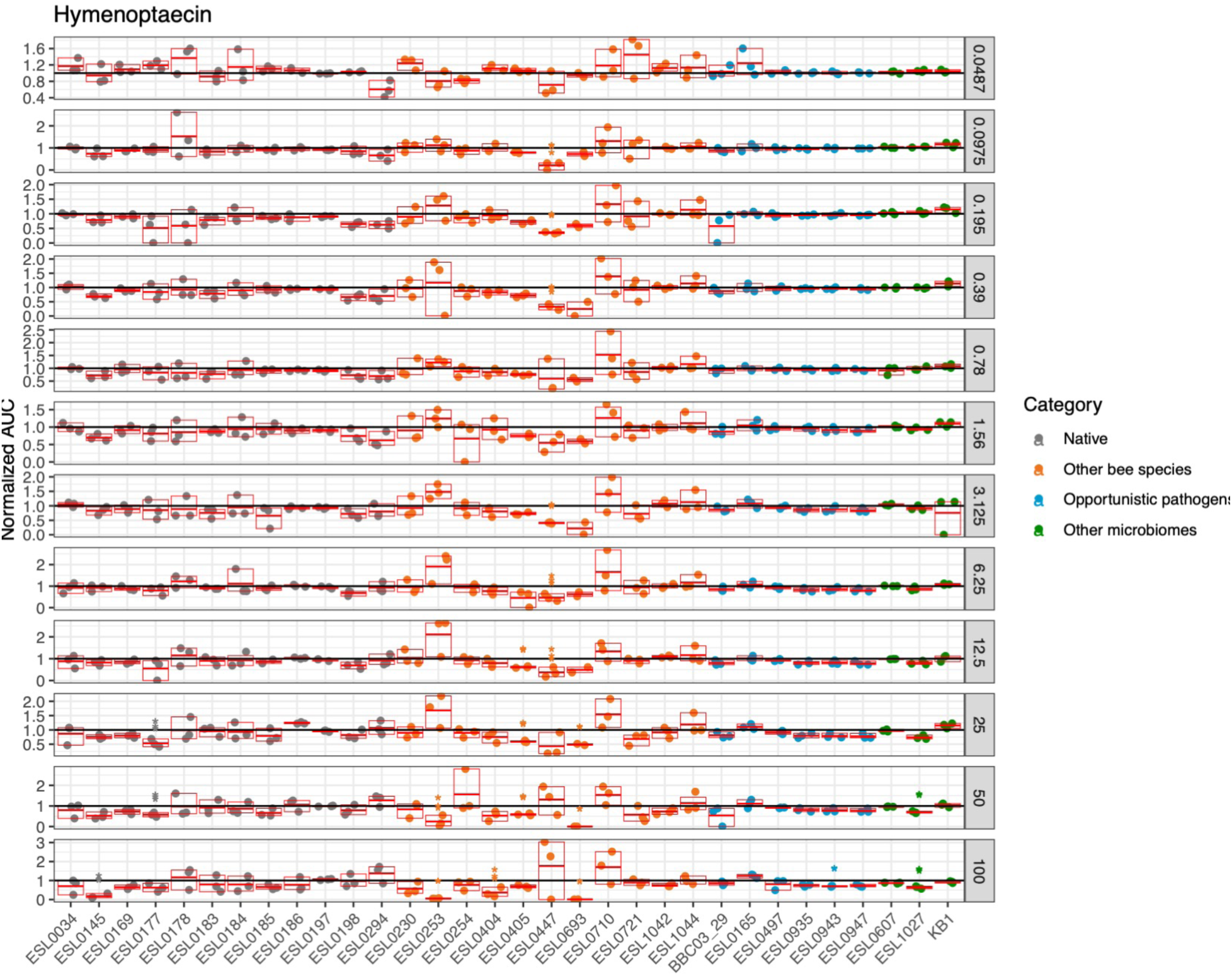
Area under the curve (AUC) of bacterial growth, normalized to the untreated control and presented on a logarithmic scale. The right panel shows the AUC values at a representative concentration of Hymenoptaecin. Each condition was tested in triplicate (n = 3). Statistical significance was assessed using a one-sample Wilcoxon test against the null hypothesis of no growth inhibition (normalized value = 1), with p-values adjusted for multiple comparisons using the false discovery rate (FDR) correction. Significance levels are indicated as follows: ****p ≤ 0.0001, ***p ≤ 0.001, **p ≤ 0.01, *p ≤ 0.05.

**Supplementary File 1**: Strain list.

**Supplementary File 2**: Primer list.

**Supplementary File 3**: qPCR abundances data.

**Supplementary File 4**: KEGG modules. Sheet 1: All KEEG modules detected. Sheet 2: Number of modules per strain with a >80% completeness, Sheet 3: Mix-model significant results with abundance data. Sheet 4: Mix-model significant results with percent of colonization data.

**Supplementary File 5**: qPCR host expression data.

**Supplementary File 6**: AMP resistance assay. Sheet 1: AMP sequence information. Sheet 2: Growth data parameters. Sheet 3: AMP sensitivity score.

**Supplementary File 7:** Genome assembly statistics for the bacterial strains sequenced in this study.

